# Robust prediction of patient outcomes with immune checkpoint blockade therapy for cancer using common clinical, pathologic, and genomic features

**DOI:** 10.1101/2023.07.04.547697

**Authors:** Tia-Gen Chang, Yingying Cao, Hannah J. Sfreddo, Saugato Rahman Dhruba, Se-Hoon Lee, Cristina Valero, Seong-Keun Yoo, Diego Chowell, Luc G. T. Morris, Eytan Ruppin

## Abstract

Despite the revolutionary impact of immune checkpoint blockade (ICB) in cancer treatment, accurately predicting patients’ responses remains elusive. We analyzed eight cohorts of 2881 ICB-treated patients across 18 solid tumor types, the largest dataset to date, examining diverse clinical, pathologic, and genomic features. We developed the LOgistic Regression-based Immunotherapy-response Score (LORIS) using a transparent, compact 6-feature logistic regression model. LORIS outperforms previous signatures in ICB response prediction and can identify responsive patients, even those with low tumor mutational burden or tumor PD-L1 expression. Importantly, LORIS consistently predicts both objective responses and short-term and long-term survival across most cancer types. Moreover, LORIS showcases a near-monotonic relationship with ICB response probability and patient survival, enabling more precise patient stratification across the board. As our method is accurate, interpretable, and only utilizes a few readily measurable features, we anticipate it will help improve clinical decision-making practices in precision medicine to maximize patient benefit.

## Introduction

Immune checkpoint blockade (ICB) has revolutionized our approach to cancer treatment in the past decade with less toxicity, durable response, and longer survival than standard therapies for patients with many types of cancer. However, many patients do not respond to these therapies, creating a need to identify biomarkers to predict which patients may derive benefit from treatment ^1–3^. Although tumor mutational burden (TMB) has been recognized as a biomarker to predict ICB efficacy in solid tumors ^4, 5^, current evidence fails to support the use of high TMB (with FDA-approved threshold of 10 Mut/Mb) as a biomarker for response to ICB treatment universally, across all cancer types ^6^. Other clinical, pathologic, and genomic features that have been reported to be associated with ICB response include tumor PD-L1 expression ^7^, microsatellite instability (MSI) ^8–10^, HLA-I evolutionary divergence (HED) ^11^, loss-of-heterozygosity status in HLA-I (LOH in HLA-I) ^12^, fraction of copy number alteration (FCNA) or tumor aneuploidy ^13, 14^, blood neutrophil-lymphocyte ratio (NLR) ^15, 16^, blood albumin level ^17^, body mass index (BMI) ^18^, sex ^19^, and age ^20^. Nonetheless, there remains an unmet need to identify factors for patient selection that are as readily measurable and provide more robust and accurate predictions for cancer ICB response and patient stratification than the approved TMB biomarker.

There have been a few attempts to integrate features from multi-omics data into a single machine-learning model to improve the predictive power of ICB response. For instance, Litchfield et al. curated 55 unique biomarkers from literature and used a tree-based ensemble model to predict ICB response with 11 most predictive biomarkers ^21^. However, this approach relied on whole-exome and transcriptome sequencing, which are expensive and not routinely measured in clinical settings. Alternatively, Chowell et al. developed a random forest model that predicts pan-cancer ICB response from 16 genomic and clinical features ^22^. The model was carefully evaluated on an independent unseen data set from the same medical center (MSK), leaving open the challenge of additional testing on external and independent cohorts. In addition, the “black box” nature of these models has limited their interpretability, impeding their application in the clinic so far.

Since cancer drug response is a complex phenomenon, it is currently challenging to perfectly distinguish responders from non-responders. Therefore, assessing the response probability of a patient to a particular therapy is of great value, potentially allowing clinicians to make more precise treatment decisions. For instance, in a patient with a high probability of ICB response, this therapy might be prioritized over another therapy; in a patient with a lower probability of ICB response, other therapeutic avenues might be prioritized. While TMB and PD-L1 expression are the two major FDA-approved biomarkers for ICB therapy, they do have limitations in accurately predicting response. For example, patients with low TMB may have a similar or even higher probability of responding to ICB therapy compared to those with high TMB ^6, 23^. Similarly, although PD-L1 expression levels are associated with response to immunotherapy in solid cancers, tumors across all PD-L1 expression levels may respond to ICB treatments ^3, 24^. Unfortunately, there has been little progress in developing a scoring system that can predict the patient-level ICB response probability in a monotonic manner, whereby higher scores reliably correlate with higher response probabilities across the entire score range.

To overcome the above-mentioned challenges, here our goal is to build a transparent white-box model to generate a reliable ICB biomarker with a few clinically easy-to-measure features, which can help clinicians to determine the patient’s probability of responding to ICB therapy. To accomplish this goal, we curated and comprehensively analyzed a large collection of pan-cancer patients, with more than 20 clinical, pathologic, and genomic features measured. We tested 20 machine-learning models to identify the most predictive model for ICB response and found that a 6-feature logistic regression model has a superior and robust performance in predicting ICB objective response on cross-validation and external test. Notably, this model identifies low TMB pan-cancer patients who can still experience benefit from immunotherapy. Simultaneously, this model demonstrates consistent predictive capabilities for response and patient survival after ICB treatment, both clinically important on their own, across almost all individual cancer types. Most importantly, the patient-level scores exhibit an almost monotonic relationship with both the ICB response probabilities, which span from 0% to 100%, and the patient survival outcomes post-treatment. To further validate the robustness and superiority of this framework, we repeated the analyses in non-small cell lung cancer (NSCLC), the cancer indication with the largest sample size in our study. Our findings suggest this approach can be a powerful tool for predicting patient clinical outcomes in ICB therapy.

## Results

### Overview

We compiled a dataset of 2,881 patients with ICB treatment across 18 solid tumor types from multiple data sources (**Fig. 1A**, **Table 1**). All patients were treated with PD-1/PD-L1 inhibitors, CTLA-4 blockade, or a combination of both immunotherapy agents (**Fig. 1A**, **Table 1**), and all samples were collected before ICB treatment.

**Figure 1.**
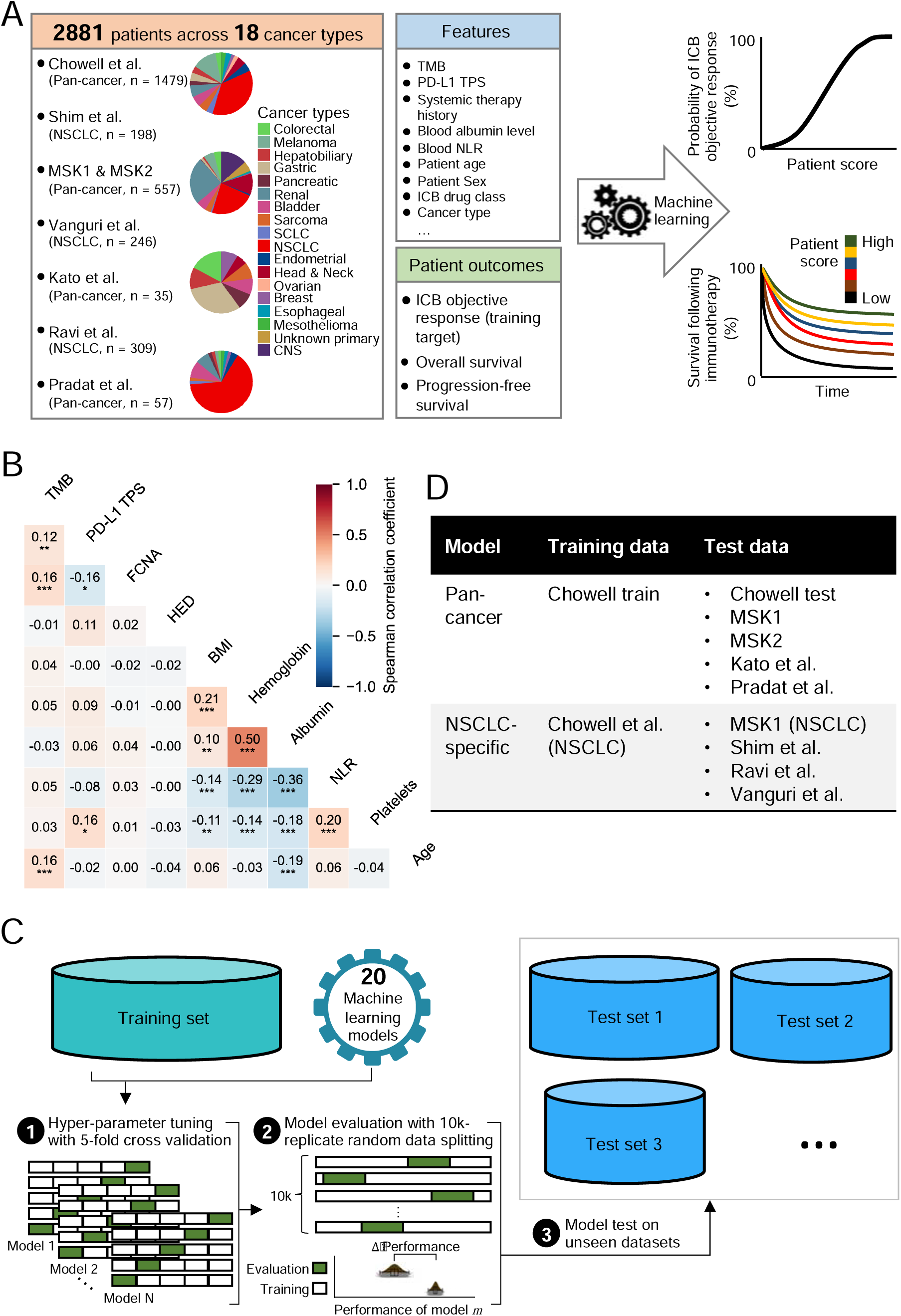
Overview of the study. **A.** Description of the study aims and data used. The study aims to develop and validate machine learning models to predict patient objective response probability and survival benefit following immunotherapy. **B.** Correlation between features measured on a continuous scale at pan-cancer level. P values and correlation coefficients are from Spearman’s rank test, and p values are further adjusted for Bonferroni correction. **C.** Schematic representation of the training, validation, and independent testing procedures used to develop and evaluate the predictive models. For each machine learning architecture, the hyperparameter is tuned with 5-fold cross-validation. After determination of the hyperparameters, the models are evaluated using various performance metrics with 2000-repeated 5-fold cross-validation. Finally, the selected models are tested on multiple unseen test cohorts to assess their generalizability. **D.** The two types of models we built, i.e., the pan-cancer and the non-small cell lung cancer (NSCLC)-specific models, and the corresponding training and test data used.

**Table 1.** Characteristics of patients in the study.

| Characteristic | Total patients | *Chowell et al. cohort* | *Shim et al. cohort* | *MSK1 cohort* | *MSK2 cohort* | *Vanguri et al. cohort* | *Kato et al. cohort* | *Ravi et al. cohort* | *Pradat et al. cohort* |
| --- | --- | --- | --- | --- | --- | --- | --- | --- | --- |
| **Sex, n (%)** |  |  |  |  |  |  |  |  |  |
| Female | 1280 (44.4) | 668 (45.2) | 58 (29.3) | 172 (38.0) | 42 (40.4) | 134 (54.5) | 17 (48.6) | 165 (53.3) | 24 (42.1) |
| Male | 1601 (55.6) | 811 (54.8) | 140 (70.7) | 281 (62.0) | 62 (59.6) | 112 (45.5) | 18 (51.4) | 144 (46.6) | 33 (57.9) |
| **Age, median, years (IQR)** | 63 (55-71) | 64 (55-71) | 62 (55-69) | 63 (53-73) | 53 (48-63) | 68 (61-73) | 62 (51-72) | 64 (57-71) | 66 (54-69) |
| **Cancer type, n (%)** |  |  |  |  |  |  |  |  |  |
| NSCLC | 1456 (50.5) | 538 (36.4) | 198 (100) | 128 (28.3) | - | 246 (100) | - | 309 (100) | 37 (64.9) |
| Renal | 232 (8.1) | 91 (6.2) | - | 137 (30.2) | - | - | - | - | 4 (7.0) |
| Melanoma | 217 (7.5) | 186 (12.6) | - | 30 (6.6) | - | - | - | - | 1 (1.8) |
| Head and neck | 132 (4.6) | 69 (4.7) | - | 61 (13.5) | - | - | 2 (5.7) | - | - |
| Bladder | 119 (4.1) | 82 (5.5) | - | 29 (6.4) | - | - | 3 (8.6) | - | 5 (8.8) |
| Sarcoma | 88 (3.1) | 67 (4.5) | - | 17 (3.8) | - | - | 3 (8.6) | - | 1 (1.8) |
| Gastric | 82 (2.8) | 64 (4.3) | - | 7 (1.5) | - | - | 11 (31.4) | - | - |
| Central nervous system | 75 (2.6) | - | - | - | 75 (72.1) | - | - | - | - |
| Colorectal | 75 (2.6) | 46 (3.1) | - | 22 (4.9) | - | - | 6 (17.1) | - | 1 (1.8) |
| Endometrial | 71 (2.5) | 65 (4.4) | - | 4 (0.9) | - | - | - | - | 2 (3.5) |
| Hepatobiliary | 62 (2.2) | 52 (3.5) | - | 5 (1.1) | - | - | 4 (11.4) | - | 1 (1.8) |
| SCLC | 55 (1.9) | 50 (3.4) | - | 4 (0.9) | - | - | - | - | 1 (1.8) |
| Esophageal | 50 (1.7) | 44 (3.0) | - | 5 (1.1) | - | - | - | - | 1 (1.8) |
| Pancreatic | 40 (1.4) | 35 (2.4) | - | 1 (0.2) | - | - | 3 (8.6) | - | 1 (1.8) |
| Mesothelioma | 36 (1.2) | 34 (2.3) | - | 1(0.2) | - | - | - | - | 1 (1.8) |
| Ovarian | 31 (1.1) | 31 (2.1) | - | - | - | - | - | - | - |
| Breast | 31 (1.1) | 25 (1.7) | - | 2 (0.4) | - | - | 3 (8.6) | - | 1 (1.8) |
| Unknown primary | 29 (1.0) | - | - | - | 29 (27.9) | - | - | - | - |
| **Drug class, n (%)** |  |  |  |  |  |  |  |  |  |
| PD-1/PD-L1 | 2447 (86.0) | 1221 (82.6) | 198 (100) | 390 (86.1) | 102 (98.1) | 234 (95.1) | - | 245 (79.3) | 57 (100) |
| CTLA-4 | 7 (0.2) | 5 (0.3) | - | 2 (0.4) | - | - | - | - | - |
| Combo | 392 (13.8) | 253 (17.1) | - | 61 (13.5) | 2 (1.9) | 12 (4.9) | - | 64 (20.7) | - |
| **Systemic therapy prior ICB, n (%)** |  |  |  |  |  |  |  |  |  |
| No | 814 (28.3) | 463 (31.3) | 14 (7.1) | 107 (23.6) | 24 (24) | 78 (31.7) | - | 123 (39.8) | 5 (8.8) |
| Yes | 2063 (71.7) | 1016 (68.7) | 184 (92.9) | 346 (76.4) | 76 (76) | 168 (68.3) | 35 (100) | 186 (60.2) | 52 (91.2) |
| **TMB, median, Mut/Mb (IQR)** | 5.9 (3.0-10.8) | 5.3 (2.8-10.8) | 7.3 (3.7-12.0) | 5.3 (3.3-8.9) | 3.9 (2.5-5.3) | 7.9 (4.4-12.3) | 7 (5-11) | 7.4 (3.4-12.7) | 7.1 (1.8-12.2) |
| **Albumin, median, g/dL (IQR)** | 3.9 (3.6-4.2) | 3.9 (3.6-4.1) | 4.1 (3.8-4.4) | 3.9 (3.6-4.2) | 4.1 (3.8-4.3) | 3.8 (3.4-4.1) | - | - | 3.9 (3.6-4.1) |
| **NLR, median, (IQR)** | 4.3 (2.7-7.0) | 4.4 (2.8-7.2) | 3.1 (1.9-4.8) | 4.1 (2.7-7.1) | 4.1 (2.5-6.8) | 4.9 (3.3-7.9) | - | - | 4.5 (2.9-6.3) |
| **PD-L1 TPS, median, % (IQR)** | 5 (0-66) | 0 (0-60) | 50 (1-72.5) | 1 (0-50) | 0 (0-50) | 5 (0-60) | - | 25 (0-75) | - |
| **ICB response, n (%)** |  |  |  |  |  |  |  |  |  |
| Responder | 825 (28.6) | 409 (27.7) | 61 (30.8) | 116 (25.6) | 14 (13.5) | 61 (24.8) | 5 (14.3) | 121 (39.2) | 19 (33.3) |
| Non-responder | 2056 (71.4) | 1070 (72.3) | 137 (69.2) | 337 (74.4) | 90 (86.5) | 185 (75.2) | 30 (85.7) | 188 (60.8) | 38 (66.7) |

The first data source was an MSK-IMPACT cohort, which included 1,479 patients treated for 16 cancer types at Memorial Sloan Kettering Cancer Center (*Chowell et al.* cohort). Sixteen clinical, pathologic, and genomic features for this cohort were reported recently ^22^. Additional information about patient PD-L1 tumor proportion score (TPS; which was available for a subset of patients), the start year of receiving ICB therapy, and the panels used for TMB determination were extracted from the electronic health records for the purpose of this study and had not been reported before. The second data source was a cohort from South Korea, which included 198 advanced NSCLC patients (*Shim et al.* cohort). Eleven features for this cohort were reported previously ^25^. Additional information about patient’s systemic therapy history before ICB therapy, blood albumin level, and blood NLR were extracted from the electronic health records for the purpose of this study and had not been reported before. The third data source was an additional cohort from Memorial Sloan Kettering Cancer Center, including 453 patients with 15 cancer types (*MSK1* cohort), and 104 patients with either central nervous system tumors or cancer of unknown primary (*MSK2* cohort). Both cohorts included 10 features. The fourth data source was a pan-cancer study from the UCSD Moores Cancer Center, consisted of 35 patients across eight cancer types (matching those in the *Chowell et al.* cohort) and six features (*Kato et al.* cohort) ^26^.

We also utilized data from the *Vanguri et al.* cohort, consisting of 246 advanced NSCLC patients at Memorial Sloan Kettering Cancer Center, with 14 available features ^27^; the Stand Up To Cancer-Mark Foundation cohort, which included 309 NSCLC patients and 10 features (*Ravi et al.* cohort) ^28^; and a pan-cancer cohort of refractory metastatic tumors (META-PRISM, *Pradat et al.* cohort) with 57 ICB-treated patients from 13 cancer types present in the *Chowell et al.* cohort and 9 features ^29^. These very recently published studies were only accessed and analyzed after our model training and initial testing had been completed and fixed.

To assess patient outcomes, three metrics were measured for all patients: objective response, progression-free survival (PFS), and overall survival (OS). Objective response was evaluated using the Response Evaluation Criteria in Solid Tumors (RECIST) ^30^ and classified as complete response or partial response for responders and stable disease or progressive disease for non-responders. Specifically, out of the total patients analyzed across various types of cancers, 825 (∼29%) experienced objective response to ICB treatment while 2,056 (∼71%) did not (**Table 1**). Across multiple cohorts, we evaluated more than 20 clinical, pathologic, and genomic features (see **Methods**). Among these features, eight were measured for most patients: patient sex, age, cancer type, ICB drug class, patient systemic therapy history before ICB treatment (whether received or not), TMB, blood albumin level, and blood NLR (**Table 1**). Additionally, the PD-L1 TPS was assessed in many NSCLC samples and a small portion of other cancer types.

We first explored the correlation between features measured on a continuous scale at pan-cancer level across all 2,881 patients (**Fig. 1B**). We found that TMB is positively correlated with FCNA (r = 0.16, adj. p < 0.001) and age (r = 0.16, adj. p < 0.001). We also observed that PD-L1 TPS is positively correlated with blood platelets (r = 0.16, adj. p < 0.05), which aligns with previous studies in ovarian cancer where platelets increase the expression of PD-L1 in tumors ^31^. Interestingly, we found that PD-L1 TPS is negatively correlated with FCNA (r = -0.16, adj. p < 0.05). In addition, a strong positive correlation was found between blood hemoglobin and albumin level (r = 0.50, adj. p < 0.001).

Benefiting from our extensive pan-cancer patient collection, complete with a comprehensive array of clinical, pathologic, and genomic features, our goal is to build a reliable ICB response predictor based on these features. The response to ICB is assessed through two main criteria: objective response, which categorizes individuals as responders or non-responders, and survival benefit, measured by PFS and OS. To achieve this, we comprehensively built and evaluated response predictors using 20 different machine learning architectures. For each model, we first tuned the optimal hyper-parameters using 5-fold cross-validation on the training set; we then evaluated its performance using 2,000-repeated 5-fold cross-validation to ensure unbiased results (equating to 10,000 random train-validation splits in total); finally, the selected models were further tested on multiple unseen test cohorts (as illustrated in **Fig. 1C**).

Our study includes two types of models: *pan-cancer* and *cancer-type-specific* (**Fig. 1D**). Pan-cancer models were developed, trained, evaluated, and compared using a subset of 964 patients from the *Chowell et al.* cohort who received immunotherapy between 2015 and 2017 (*Chowell training*). The unseen test cohorts include 515 patients from the *Chowell et al.* cohort who received immunotherapy in 2018 (*Chowell test*), as well as patients from the *MSK1*, *MSK2*, *Kato et al.*, and *Pradat et al.* cohorts. Cancer-type-specific models were developed, trained, evaluated, and compared using the *Chowell et al.* cohort (NSCLC patients only), unseen test cohorts include patients from the *MSK1* (NSCLC patients only), *Shim et al.*, *Ravi et al.*, and *Vanguri et al.* cohorts. This approach allowed us to thoroughly evaluate the generalizability of both our pan-cancer and NSCLC-specific models across multiple independent datasets.

### Robust prediction of pan-cancer objective response to immunotherapy by a 6-variable logistic LASSO regression model

We first developed a logistic regression model to predict the pan-cancer objective response to ICB therapy using the 8 features shared among all patients. We performed 5-fold cross-validation to identify the optimal hyperparameters of the model. Interestingly, our analysis showed that the patient’s sex and ICB drug class information had little impact on the prediction (**SFig. 1A**), and thus those were excluded from the final model. After hyperparameter tuning, the best model found is a 6-feature logistic LASSO regression model (LLR6), which includes the following features in decreasing order of importance: TMB, systemic therapy history, blood albumin, blood NLR, age, and cancer type (**SFig. 1B**). Here *systemic therapy history* is a binary variable and indicates whether the patient received chemotherapy or targeted therapy prior to immunotherapy.

To double-check and assess the possible added value of the other features, we also developed and tuned a logistic regression model using all 16 features. However, the 16-feature model showed no significant improvement over LLR6 on the cross-validation sets (**SFig. 2**). In addition, we tested a five-variable logistic regression model without using TMB, but it performed worse than LLR6 (**SFig. 2**). Overall, our analysis suggests that the selected 6 features capture the best essential information for predicting ICB response in pan-cancer patients.

To further test if there were better machine learning architectures for predicting ICB response using all 16 features, we experimented with 15 additional machine learning models, such as decision trees, Gaussian processes, support vector machine, XGBoost, and deep neural networks, among others. However, none of these models outperformed LLR6 on the cross-validation sets. While some complex models, such as XGBoost and a 2-layer deep neural network, showed comparable performance to LLR6, they exhibited much larger discrepancies in performance between the training and validation data. This suggests a high risk of overfitting the data (**SFig. 2**).

To further compare the performance of LLR6 with established methods, we evaluated its predictive power against the 16-feature random forest model (RF16) reported by Chowell et al. ^22^ and the FDA-approved TMB biomarker. In addition, we constructed a 6-feature random forest model (RF6) and optimized its hyperparameters using the protocol described by Chowell et al. ^22^. Firstly, we found that LLR6 outperformed the TMB biomarker, demonstrating a synergistic effect in predicting ICB response by integrating multiple features (**Fig. 2A**). Moreover, LLR6 also outperformed both RF16 and RF6 (**Fig. 2A**). Notably, LLR6 exhibited a close-to-zero performance difference between training and cross-validation, which is much smaller than that of RF16 and RF6 models and even comparable to the unsupervised TMB biomarker (**Fig. 2B**), which suggests that LLR6 is less prone to overfitting.

**Figure 2.**
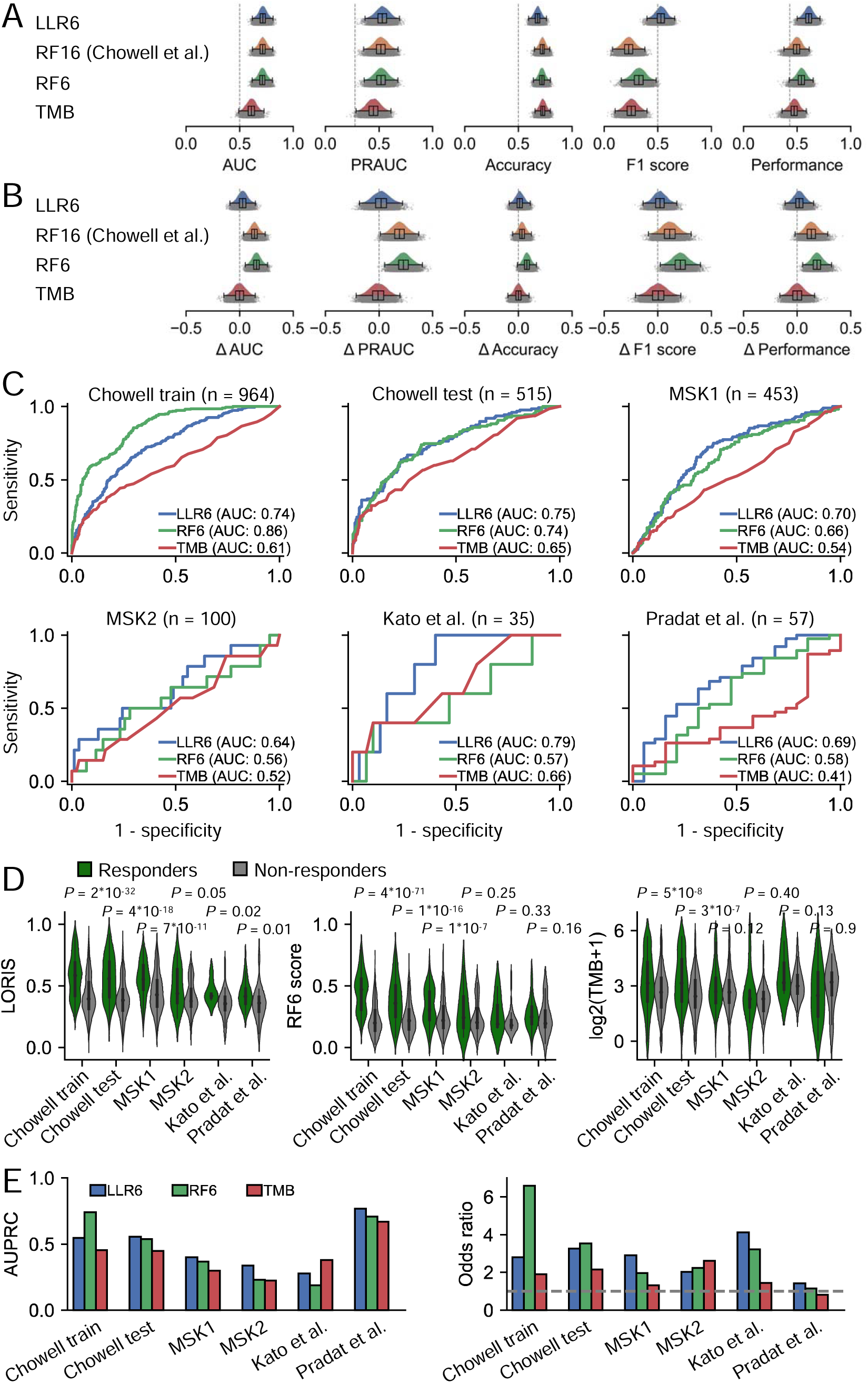
Robust prediction of pan-cancer objective response to immunotherapy by a 6-variable logistic LASSO regression model. **A-B.** Comparison of model predictive power on cross-validation sets (**A**) and the difference in model predictive power between training and cross-validation (**B**) for LLR6, RF16 (Chowell et al.), RF6, and the TMB biomarker. Here *performance* is defined as the geometric mean of the area under the receiver operating characteristic curve (AUC), the area under the precision-recall curve (AUPRC), the accuracy, and the F1 score. **C.** Receiver operating characteristic curves and corresponding AUC of LLR6, RF6, and the TMB biomarker on the training set and across multiple unseen test sets. **D.** Distribution of Logistic Regression-based Immunotherapy-response Score (LORIS), RF6 score, and TMB alone in responders and non-responders on the training set and across multiple unseen test sets. Two-sided p-values were calculated using the Mann-Whitney U test. The upper and lower boundaries signify the first and third quartiles, correspondingly, while the central line denotes the median. Whiskers stretch to the most distant data points not classified as outliers (within 1.5 times the interquartile range), and outliers are illustrated as points above and below the box-and-whisker diagram. **E.** AUPRC and odds ratio of ICB objective response of LLR6, RF6, and the TMB biomarker on the training set and across multiple unseen test sets.

To directly assess the generalizability of LLR6, we applied it to five unseen test datasets. The regression coefficients (including the intercept) of LLR6 used for prediction purposes are the average corresponding values obtained in the previously described 10,000 training iterations without any further adaptation on the test data. We referred to the output calculated via LLR6 as the Logistic Regression-based Immunotherapy-response Score (LORIS), calculated using the formula described in the **Methods** section.

LLR6 consistently outperformed RF6 and TMB biomarker across all test sets, even for cancer types not seen in the training data such as central nervous system tumors and cancer of unknown primary in the *MSK2* cohort. Specifically, LLR6 achieved higher area under the receiver operating characteristic curve (AUC) on the *Chowell test* (LLR6 AUC = 0.75, RF6 AUC = 0.74, TMB AUC = 0.65), *MSK1* (LLR6 AUC = 0.70, RF6 AUC = 0.66, TMB AUC = 0.54), *MSK2* (LLR6 AUC = 0.64, RF6 AUC = 0.56, TMB AUC = 0.52), *Kato et al.* datasets (LLR6 AUC = 0.79, RF6 AUC = 0.57, TMB AUC = 0.66), and *Pradat et al.* datasets (LLR6 AUC = 0.69, RF6 AUC = 0.58, TMB AUC = 0.41) (**Fig. 2C**). Simultaneously, LLR6 consistently outperformed RF6 and the TMB biomarker by predicting significantly higher LORIS for responders compared to non-responders. (**Fig. 2D**). In addition, LLR6 showed a superior area under the precision-recall curve (AUPRC) on most datasets compared to RF6 and TMB biomarker (**Fig. 2E**).

To binarize the values of LORIS and RF6 scores, we used cutoffs of 0.5 and 0.27 respectively, which maximize sensitivity and specificity of ICB response prediction on the training data. Regarding TMB, the FDA-approved cutoff of 10 Mut/Mb was used. As a result, LLR6 performed similarly to or better than RF6 and the TMB biomarker in terms of various metrics calculated using binarized scores, including accuracy, F1 score, positive predictive value (PPV), negative predictive value (NPV), specificity, and sensitivity (**SFig. 3**). In particular, LLR6 predicted an odds ratio of 1.4 - 4.1 for ICB objective response between high-LORIS and low-LORIS patients across different test sets, which is in general higher than RF6 (1.2 - 3.5) and the TMB biomarker (0.8 - 2.6) (**Fig. 2E**).

### LORIS successfully identifies low TMB patients who can benefit from immunotherapy

We conducted further investigation to determine whether LORIS could be useful in stratifying patients by their survival outcomes following immunotherapy. To ensure we had a sufficiently large sample size for each cancer type, we combined patients from two test sets (*Chowell test* and *MSK1*; n = 968), which includes all cancer types covered in the training data and has all six input features and three patient outcomes measured. Our pan-cancer Kaplan-Meier analysis revealed that patients with low LORIS (binned at 0.5) had significantly worse survival compared to those with high scores (OS HR = 3.2, 2*10^-28^, **Fig. 3A**; PFS HR = 2.6, p = 2*10^-33^, **SFig. 4A**). In contrast, using TMB (binned at 10 Mut/Mb) to stratify patients resulted in moderate power (OS: HR = 1.3, p = 0.01, **Fig. 3A**; PFS: HR = 1.5, p = 8*10^-6^, **SFig. 4A**). Notably, LORIS identified a substantial proportion of low TMB patients who could benefit from immunotherapy to a similar extent as high TMB patients (**Fig. 3A; SFig. 4A**). Similar results were obtained when we used the 50th percentile in each cancer type as the optimal cutoff for LORIS binning and the top 20th percentile for TMB binning, as proposed by Samstein et al. ^5^ (**Fig. 3B; SFig. 4B**).

**Figure 3.**
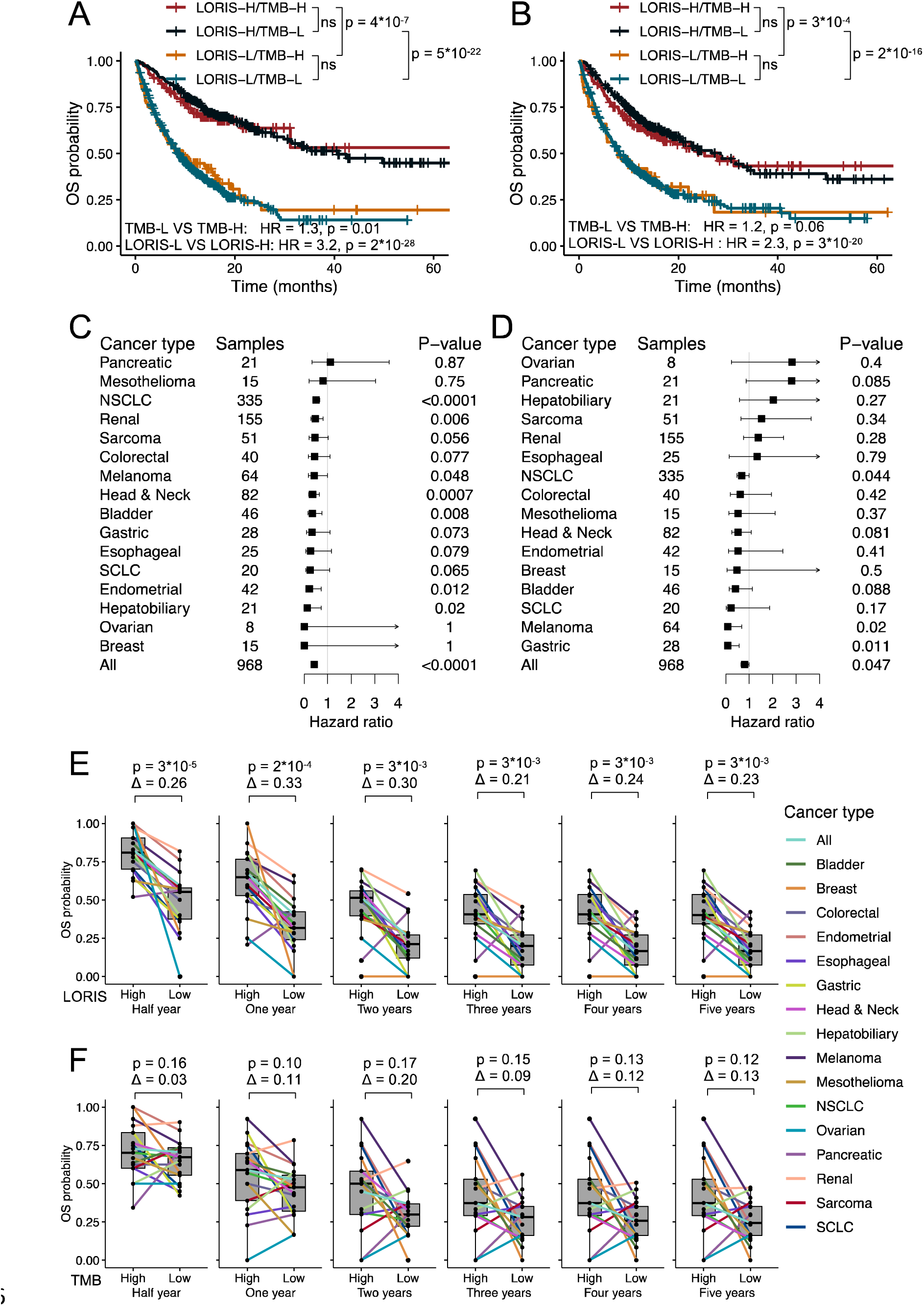
LORIS predicts patient outcomes following immunotherapy for both pan-cancer and individual cancer types. **A.** Kaplan–Meier analysis of TMB binned at 10 Mut/Mb and LORIS binned at 0.5. P values next to the legend indicate pairwise comparisons. H, high; L, low. **B.** Same as panel A, but TMB is binned at the highest 20th percentile and LORIS is binned at the 50th percentile for each cancer type. **C-D.** Forest plot of OS within each cancer type using LORIS (binned at the 50th percentile; **C**) or TMB (binned at the highest 20th percentile; **D**) in a multivariable model with adjustment for cancer type, age, ICB drug class, and year of ICB start. Squares positioned at midpoints symbolize point estimates of HRs, and the accompanying bars indicate 95% confidence intervals. **E-F.** Comparison of half-year, 1-year, 2-year, 3-year, 4-year, and 5-year OS stratified by cancer type for high versus low LORIS (binned at the 50th percentile; **E**) and high versus low TMB (binned at the highest 20th percentile; **F**). Survival probability differences and paired Wilcoxon-test p-values are displayed in panels E and F. The upper and lower boundaries signify the first and third quartiles, correspondingly, the central line denotes the median, and the whiskers stretch to the most distant data points not classified as outliers (within 1.5 times the interquartile range). Data are from combined *Chowell test* (n = 515) and *MSK1* (n = 453) sets.

To test the predictive power of LLR6 in individual cancer types, we calculated HRs for LORIS and TMB for each cancer type using a multivariate Cox regression model that accounted for age, ICB drug class, and year of ICB start. Consequently, higher LORIS predicted lower than 1 OS HRs for all except one individual cancer types (binned at the 50th percentile; **Fig. 3C**), which is not true for the TMB biomarker (binned at the top 20th percentile; **Fig. 3D**). Similar results were also observed for PFS (**SFig. 4C,D**).

We also examined whether higher LORIS could predict better short-term and/or long-term patient survival, as both metrics are clinically important on their own. We compared the survival probability between patients with high versus low LORIS at various time points after immunotherapy, including half a year, one year, two years, three years, four years, and five years. Notably, higher LORIS predicted significantly better OS for all time points (difference in survival probability between high LORIS versus low LORIS patients: 0.21 – 0.33; Wilcoxon-test p values: 3*10^-5^ – 3*10^-3^; **Fig. 3E**). In contrast, TMB did not consistently predict better OS for all time points (difference in survival probability between high TMB and low TMB patients: 0.03 – 0.20; Wilcoxon-test p values: 0.10 – 0.17; **Fig. 3F**). We also observed similar results for PFS (**SFig. 4E,F**).

### Monotonic relationship between LORIS and patient objective response probability & survival following immunotherapy

Next, we investigated the relationship between a patient’s LORIS and their response to ICB therapy. Notably, we uncovered a unique characteristic of the LORIS signature. Specifically, as the LORIS increased, there was a nearly consistent rise in the probability of objective response for patients, ranging from 0% to 100% (**Fig. 4A**). This distinctive attribute allows clinicians to estimate the odds of ICB response for cancer patients by measuring the six input features and calculating the LORIS. In particular, LORIS enables the identification of the top 10% of patients who are highly likely to respond to ICB therapy (with a response probability exceeding 50%) while excluding the bottom ∼10% of patients who are unlikely to respond (with a response probability below 10%). In contrast, the stratification power of TMB falls short. Only the top 6% of patients with the highest TMB exhibited a response probability exceeding 50%, and the lowest TMB scores proved ineffective in excluding non-responsive patients altogether (**Fig. 4B**).

**Figure 4.**
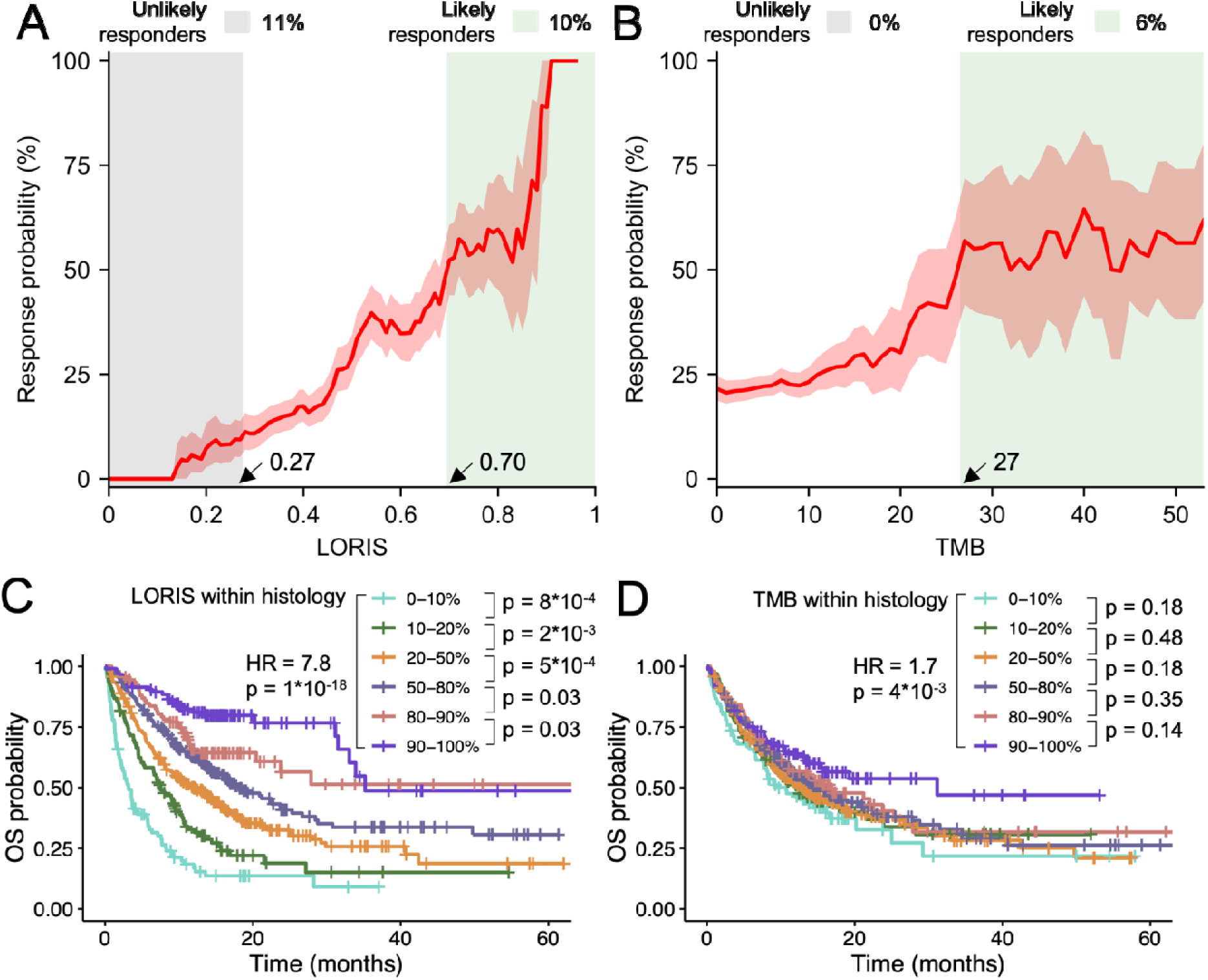
Monotonic relationship between LORIS and patient objective response probability & survival following immunotherapy. **A-B.** Relationship between LORIS (**A**) or TMB (**B**) and ICB objective response probability. The average values and 95% confidence interval are shown using 1,000-replicate bootstrapping. The grey region represents patients with an unlikely response to immunotherapy (with a response probability below 10%), while the green regions represent patients with a likely response (with a response probability exceeding 50%). The arrows indicate the LORIS and TMB threshold values. **C-D.** Kaplan–Meier analysis of LORIS (**C**) or TMB (**D**) binned at the different percentiles in each cancer type. In panels C and D, p values next to the legend indicate pairwise single-tail comparisons testing against the hypothesis that “higher scored patients do not have better survival than lower scored patients”; hazard ratios (HRs) are shown for the lowest-percentile (0-10%) and the highest-percentile groups (90-100%). Data are from combined *Chowell test* (n = 515) and *MSK1* (n = 453) sets.

We further tested whether LORIS could predict patient survival following immunotherapy. We found that higher patient LORIS consistently predicted better OS for patients across different percentiles. We were able to group patient survival into as many as six categories based on LORIS within each cancer type, i.e., 0-10% < 10-20% < 20-50% < 50-80% < 80-90% < 90-100%. Notably, the HR between the lowest (0-10%) and highest (90-100%) percentile groups was as high as almost eight (**Fig. 4C**). However, we did not observe this monotonic relationship with survival for the TMB biomarker (**Fig. 4D**). Similar results were observed for PFS (**SFig. 5**). These findings demonstrate the potential of LORIS as a robust and reliable biomarker for predicting patients’ objective response as well as survival following immunotherapy.

**Figure 5.**
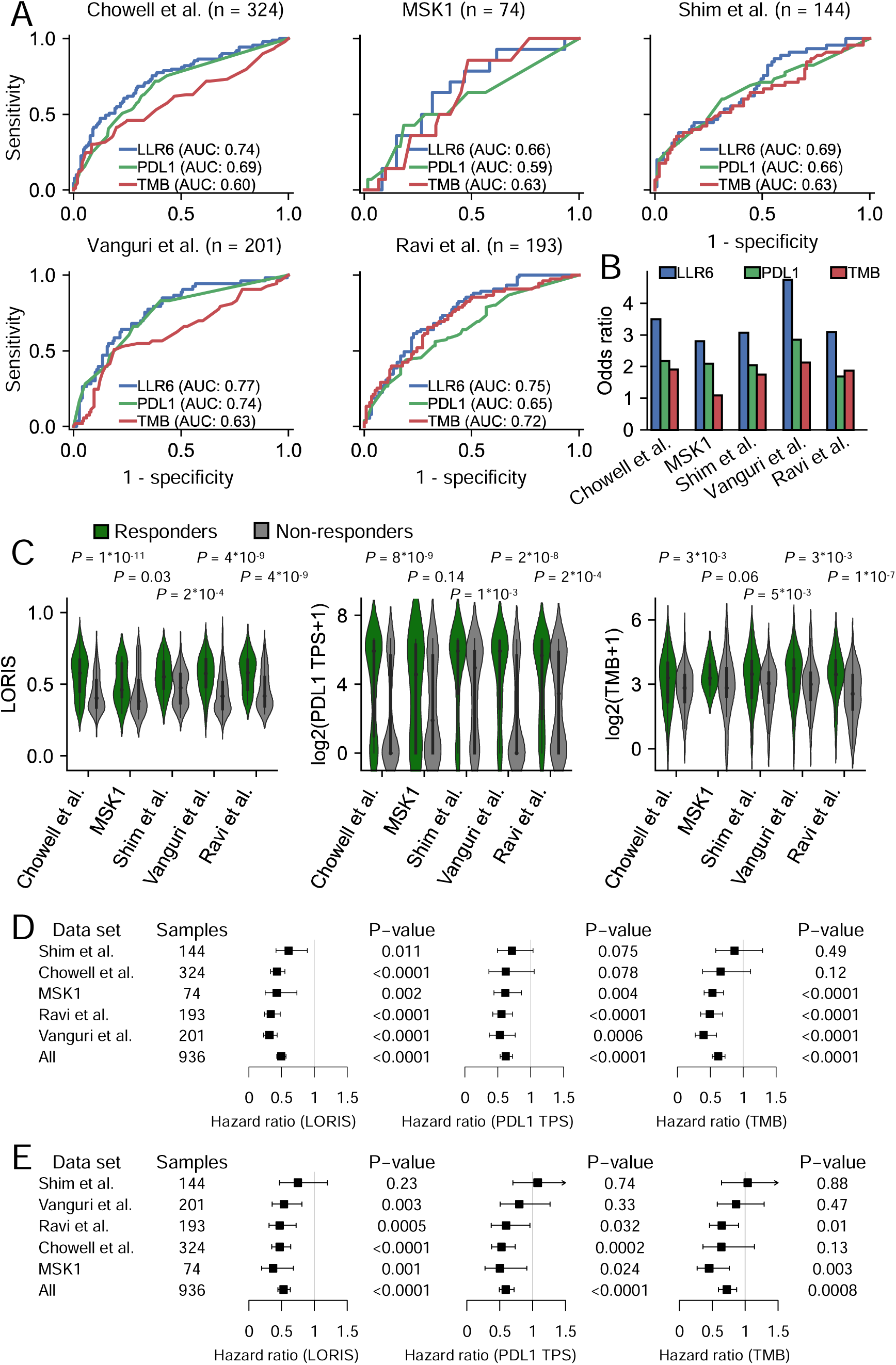
Robust prediction of response to immunotherapy in NSCLC with logistic LASSO regression. **A.** Receiver operating characteristic curves and corresponding area under the curve (AUC) of LLR6, PD-L1 TPS, and TMB on training set and across multiple unseen test sets. **B.** Odds ratio of ICB objective response of LLR6, PD-L1 TPS, and the TMB biomarker on the training set and across multiple unseen test sets. **C.** Distribution of LORIS, PD-L1 TPS, and TMB in responders and non-responders on training set and across multiple unseen test sets, with P values calculated using the Mann–Whitney U test. The upper and lower boundaries signify the first and third quartiles, correspondingly, while the central line denotes the median. Whiskers stretch to the most distant data points not classified as outliers (within 1.5 times the interquartile range), and outliers are illustrated as points above and below the box-and-whisker diagram. **D-E.** Forest plots of PFS (**D**) and OS (**E**) within each data set using LORIS (binned at 0.44, which maximize sensitivity and specificity of ICB response prediction on the training data), or PD-L1 TPS (binned at 50%), or TMB (binned at 10 Mut/Mb) in a multivariable model with adjustment for age and ICB drug class. Squares positioned at midpoints symbolize point estimates of HRs, and the accompanying bars indicate 95% confidence intervals.

### Robust prediction of cancer-type-specific response to immunotherapy with logistic LASSO regression

Our study has demonstrated the superior capability of our pan-cancer model in integrating diverse clinical variables for precise patient stratification and improved prediction of ICB response across various cancer types. This naturally led us to probe a subsequent question: could the approach proposed herein be successfully employed to develop cancer-type-specific models? The rationale behind posing this question lies in the potential enhancement of predictive power that might be realized if a model can incorporate features specific to a particular type of cancer, provided sufficient training data is available. In pursuit of this goal, we investigated the potential of employing logistic LASSO regression to create a specific model for NSCLC, the cancer type with the largest sample size in our dataset.

We constructed, trained, and assessed NSCLC-specific models utilizing a similar protocol to that used in our pan-cancer study, albeit with two minor adjustments. First, we harnessed the entire *Chowell et al.* cohort as our training data to ensure we had an adequate number of samples for model training. Secondly, we replaced the cancer type feature in the pan-cancer LLR6 model with PD-L1 TPS. Given that we were dealing with only one type of cancer in this instance, the cancer type feature carried no meaningful information. Furthermore, PD-L1 TPS is one of the most important biomarkers in NSCLC and is routinely measured for the majority of NSCLC patients during clinical practice. Consequently, a total of 324 NSCLC patients in the training dataset were evaluated with all six input features for LLR6, specifically TMB, PD-L1 TPS, patient’s systemic therapy history, albumin, NLR, and patient age.

As a result, the NSCLC-specific LLR6 model was among the best models with 2000-repeated 5-fold cross-validation compare with other 19 different models. More importantly, despite the limited data size, it maintained a near-zero performance discrepancy between the training and cross-validation data. This suggests a minimized risk of overfitting and enhanced robustness in predicting ICB response (**SFigs. 6**). Indeed, NSCLC-specific LLR6 consistently outperformed both the TMB biomarker and the PD-L1 TPS biomarker in predicting ICB objective response on all five data sets (LLR6: AUC = 0.66 – 0.77; PD-L1 TPS: AUC = 0.59 – 0.74; TMB: AUC = 0.60 – 0.72; **Fig. 5A**).

Moreover, the NSCLC-specific LORIS (see the formula calculating it in **Methods**) predicted an odds ratio of 2.8 - 4.7 for ICB objective response between high-LORIS and low-LORIS patients, which is much higher than that of the TMB biomarker (1.1 - 2.1) or the PD-L1 TPS biomarker (1.7 - 2.9) (**Fig. 5B**). In addition, responders consistently had significantly higher LORIS than non-responders across all data sets (**Fig. 5C**). Lastly, higher LORIS consistently predicted lower risks (HRs < 1) for both PFS and OS on all datasets, after adjusting for age and ICB drug class (**Fig. 5D,E**). These findings suggest that our approach can indeed be extended to construct robust, cancer-type-specific models.

## Discussion

The clinical utility of many machine learning models is markedly hindered by their “black box” nature, which makes them difficult to interpret ^32–34^. To address this issue, here we developed two transparent and interpretable logistic regression models, one applicable pan-cancer and one for NSCLC patients, that integrate six commonly measured clinical features to predict patient ICB response. Our results demonstrate three key findings. Firstly, LORIS, the score predicted by our models, not only predicts patient ICB objective response but also predicts short-term and long-term survival benefit following immunotherapy better than the existing methods. Secondly, and most importantly, our model successfully identifies low TMB or low PD-L1 TPS patients who can still benefit from immunotherapy. Finally, compared to the clinically used TMB and PD-L1 TPS biomarkers, LORIS scores patients by their response probabilities to immunotherapy in a much more monotonic and consistent manner, resulting in more accurate identification of highly likely responders and exclusion of likely non-responders. Our findings suggest that LORIS could be a valuable tool for improving clinical decision-making practices in precision medicine to maximize patient benefit.

We have identified key features that can effectively predict a patient’s ICB response, including TMB, systemic therapy history, albumin, NLR, age, cancer type (specifically for the pan-cancer model), and PD-L1 TPS (specifically for the NSCLC-specific model). Notably, a patient’s systemic therapy history, while not typically considered in ICB response prediction, plays a significant role in both models. Since immunotherapies activate and amplify the immune system, individual immune competency and diversity can influence differential response rates; theoretically, chemotherapy reduces immune system competency and could lead to reduced ICB response rates ^35^. Indeed, He et al. showed that first-line chemotherapy can influence the tumor microenvironment and decrease the efficacy of subsequent immunotherapy through whole-exome sequencing and RNA sequencing in 32 NSCLC patients ^36^. Furthermore, it was observed that resistance to anti-MAPK targeted therapy could promote an immune-evasive tumor microenvironment and cross-resistance to subsequent immunotherapy in melanoma cases ^37^. More recently, it was found that removing systemic therapy history decreased the predictive power of ICB response ^22^. However, we should note that a patient’s systemic therapy history is not a tumor-intrinsic feature, likely also reflects how heavily pre-treated a patient is (and may therefore be associated with poorer performance status and overall prognosis), and may be influenced by various factors that guide an oncologist’s decision to opt for chemotherapy, targeted therapy, or immunotherapy as first-line treatment. Consequently, further research is necessary to identify the critical intrinsic factors influenced by a patient’s systemic therapy history that contribute to varied ICB responses. We anticipate that a deeper understanding of these elements could further augment the predictive power of our model.

Our analyses reveal two interesting implications. First, our logistic regression model outperforms other more complex models in predicting ICB response in both pan-cancer and NSCLC-specific scenarios, suggesting that the effect of interaction between these clinical, pathologic, and genomic variables on ICB response prediction is minor, at least in our dataset. This finding is consistent with a previous study that used clinical and utilization data to predict early detection of type 2 diabetes, where logistic LASSO regression performed better than or comparably to other complex models, such as random forests, gradient-boosted decision trees, and neural networks ^38^. However, further testing of this implication is needed in other cohorts and for other features such as gene mutation signatures, transcriptomic features, and imaging features. Second, our study shows that a few clinical features are sufficient to predict ICB response, despite the availability of more than 20 features in total. This phenomenon is consistent with a recent meta-analysis of genomic and transcriptomic biomarkers of ICB response, where simple signatures involving a few genes, such as the PD-L1 signature, the CYT signature, the IFNG signature, and the CD8_Jiang signature, outperformed many complex signatures involving tens or hundreds of genes ^39^. The redundancy of information among features could explain why some features, such as blood hemoglobin and platelets, show high importance in a previous study^22^ but can be omitted by LLR6 without loss of predictive power since these features are significantly correlated with blood albumin and NLR, respectively (**Fig. 1B**). Alternatively, some features may work specifically in certain cancer types, such as MSI (or mismatch repair deficiency) in locally advanced rectal cancer ^40^, FCNA in melanoma and renal cell carcinoma ^14^, and HLA LOH and HLA divergence in several cancer types but not in NSCLC ^11, 12, 41^.

This study has a few limitations. Firstly, our study had a retrospective design, and in order to further demonstrate the transformative value of LLR6 in clinical settings, more prospective studies need to be conducted in the future. Secondly, although we curated the largest cohort with comprehensive clinical, pathologic, and genomic features measured in a single study, the sample size is still limited for most individual cancer types. As a result, we could only build cancer-type-specific models for NSCLC. Moreover, the patient population in our study may not be representative of the global cancer patient population, which could introduce both demographic and behavioral bias. However, we can easily retrain our models when more representative data becomes available. Additionally, we did not have transcriptomic data for the patients, which is an important factor in assessing tumor microenvironment and predicting ICB response ^21, 39, 42–44^. Finally, the PD-L1 TPS data was mainly limited to NSCLC patients and rarely measured in other cancer types. Despite this limitation, our preliminary analysis shows that the NSCLC-specific LLR6 model, which incorporates the PD-L1 TPS information, can also enhance the predictive power of ICB objective response in other cancer types, such as gastric cancer, esophageal cancer, and mesothelioma (gastric cancer, n = 27, LLR6 AUC = 0.72, PD-L1 AUC = 0.36, TMB AUC = 0.50; esophageal cancer, n = 15, LLR6 AUC = 0.58, PD-L1 AUC = 0.51, TMB AUC = 0.53; mesothelioma, n = 18, LLR6 AUC = 0.80, PD-L1 AUC = 0.56, TMB AUC = 0.41; **SFig. 7**). However, further validation is needed to confirm the importance of PD-L1 expression information in predicting ICB response in individual cancer types using more extensive cohorts.

In summary, this study analyzed the largest and most diverse cohort to date of immunotherapy-treated cancer patients with both clinical, pathologic, and genomic data and ICB response information, which allowed us to develop a robust machine learning compact model to predict patient’s objective response and survival following immunotherapy. Our model relies on a few easily measurable clinical features and produces monotonic scores, which have the potential to aid clinical decision-making and patient stratification. As our understanding of tumor immunology and the availability of comprehensive data in larger cohorts continues to improve, we expect to see the development of even more accurate models for personalized precision therapy and ultimately reducing cancer mortality.

## Methods

### Cohort description

All patient features in the following cohorts were collected before the start of ICB treatment.

#### Chowell et al. cohort

The *Chowell et al.* cohort comprises 1,479 patients diagnosed with 16 different types of solid tumors. The cohort data includes measurements of 19 features. Sixteen of them were previously reported, including TMB, FCNA, HED, LOH in HLA-I, MSI status, BMI, patient systemic therapy history before immunotherapy, sex, age, NLR, albumin, platelets, hemoglobin, cancer type, tumor stage, and ICB drug class. TMB is calculated as the total number of somatic non-synonymous mutations in the tumor, normalized to the exonic coverage of the respective MSK-IMPACT panels in Mut/Mb. For more information about how patient features were measured and outcomes were assessed, please refer to Chowell et al. ^22^. We downloaded deidentified data from Supplementary Table 3 of Chowell et al. ^22^. Three features, i.e., patient PD-L1 TPS (which was available for a subset of patients), the start year of receiving ICB therapy, and the panels used for TMB determination (which identifies non-synonymous mutations in 341 or 410 or 468 genes) were extracted from the electronic health records for the purpose of this study and was not reported before. PD-L1 TPS was determined using the Dako PD-L1 IHC 22C3 pharmDx kit (Agilent Technologies, Santa Clara, CA), which is approved by the US Food and Drug Administration.

#### Shim et al. cohort

The *Shim et al.* cohort includes 198 patients with advanced NSCLC, with 13 features measured. These features include tumor PD-L1 expression, TMB, LOH in HLA-I, patient systemic therapy history before immunotherapy, sex, age, NLR, albumin, smoking status (ever [former or current] vs never), histology (lung adenocarcinoma vs squamous cell carcinoma vs others), Eastern Cooperative Oncology Group (ECOG) performance status (from 0 to 2), and ICB drug class. TMB was defined as the number of nonsynonymous alterations (single nucleotide variations and indels), identified from whole-exome sequencing. PD-L1 expression was assessed using the US Food and Drug Administration-approved Dako PD-L1 IHC 22C3 pharmDx kit (Agilent Technologies, Santa Clara, CA) in the samples. Detailed information on the measurement of patient features and outcomes can be found in Shim et al. ^25^. All deidentified data were obtained from Supplementary Table 1 of Shim et al. ^25^, except for blood NLR, albumin level, and systemic therapy history, which were extracted from the electronic health records for the purpose of this study and was not reported before.

#### MSK1 & MSK2 cohorts

The use of the patient data from the *MSK1* & *MSK2* cohorts was approved by the Memorial Sloan Kettering Cancer Center institutional review board. The research question (whether integrated machine learning model could predict patient outcomes with ICB therapy) was specified before data collection began. Patients selected for this study were those with solid tumors diagnosed from 2014 through 2019 who received at least 1 dose of ICB at Memorial Sloan Kettering Cancer Center. We excluded patients with a history of more than one cancer, those without a complete blood count within 30 days prior to the first dose of ICB, those enrolled in blinded trials. We excluded patients who received ICB in a neoadjuvant or adjuvant setting, and patients with unevaluable response. The final set consisted of 557 patients with solid tumors from 17 different types. Thirteen features were measured in the study, including TMB, FCNA, patient systemic therapy history before immunotherapy, sex, age, NLR, albumin, platelets, cancer type, ICB drug class, patient PD-L1 TPS (which was available for a subset of patients), the start year of receiving ICB therapy, and the panels used for TMB determination. The measurement of clinical and genomic features is the same as that of the *Chowell et al.* cohort.

#### Vanguri et al. cohort

The *Vanguri et al.* cohort includes 247 patients with advanced NSCLC, with 15 features measured. One sample with unknown primary tumor site was excluded. These features include tumor PD-L1 expression, TMB, FCNA, MSI status, patient systemic therapy history before immunotherapy, sex, age, derived NLR (defined as ratio of neutrophils to the difference between total leukocytes and neutrophils in peripheral blood), albumin, smoking status, tobacco use, histology, ECOG performance status, ICB drug class, and the panels used for TMB determination. TMB is calculated as the total number of somatic non-synonymous mutations in the tumor, normalized to the exonic coverage of the respective MSK-IMPACT panels in Mut/Mb. PD-L1 immunohistochemistry was performed on 4-_μ_m FFPE tumor tissue sections using a standard PD-L1 antibody (E1L3N; dilution 1:100, Cell Signaling Technologies) validated in the clinical laboratory at the study institution. Detailed information on the measurement of patient features and outcomes can be found in Vanguri et al. ^27^. All deidentified data were obtained from synapse: https://www.synapse.org/#!Synapse:syn26642505 and cbioportal: https://www.cbioportal.org/study/summary?id=lung_msk_mind_2020.

#### Kato et al. cohort

The *Kato et al.* cohort comprises 429 patients, where 35 patients from eight solid tumor types were included in this study based on three criteria: (1) patients received immunotherapy, (2) their cancer types are included in the *Chowell et al.* cohort, and (3) TMB was measured. Six features were assessed, namely TMB, MSI status, patient systemic therapy history before immunotherapy, sex, age, and cancer type. TMB was determined using panel NGS performed by a CLIA-certified laboratory. More detailed information on patient feature measurement and outcomes can be found in Kato et al. ^26^. Deidentified data were obtained from Supplementary Data 1 of Kato et al. ^26^. It should be noted that blood NLR and albumin levels were unavailable for this cohort, and only four variables were used in the pan-cancer LLR6 model, i.e., TMB, patient systemic therapy history before immunotherapy, age, and cancer type.

#### Ravi et al. cohort

The *Ravi et al.* cohort includes 393 patients with NSCLC treated with anti-PD-(L)1 therapy, with 10 features measured, where 309 patients with TMB measured were included in this study. These features include tumor PD-L1 expression, TMB, patient systemic therapy history before immunotherapy, sex, age, tumor stage, smoking status, tobacco use, histology, and ICB drug class. TMB was defined as the number of nonsynonymous alterations (single nucleotide variations, de novo variants, and indels), identified from whole-exome sequencing at the Genomics Platform of the Broad Institute of Harvard and MIT. Detailed information on the measurement of patient features and outcomes can be found in *Ravi* et al. ^28^. All deidentified data were obtained from Zenodo repository: https://doi.org/10.5281/zenodo.7625517. It should be noted that blood NLR and albumin levels were unavailable for this cohort, and only four variables were used in the NSCLC-specific LLR6 model, i.e., TMB, patient systemic therapy history before immunotherapy, age, and PD-L1 TPS.

#### Pradat et al. cohort

The *Pradat et al.* cohort comprises 1031 pan-cancer patients, where 57 patients from 13 solid tumor types were included in this study based on two criteria: (1) patients received immunotherapy, (2) their cancer types are included in the *Chowell et al.* cohort, and (3) TMB was measured. Nine features were assessed, i.e., TMB, FCNA, patient systemic therapy history before immunotherapy, NLR, albumin, sex, age, ICB drug class, and cancer type. TMB was determined using whole exome sequencing. More detailed information on patient feature measurement and outcomes can be found in Pradat et al. ^29^. Deidentified data were obtained from Supplementary Tables of Pradat et al. ^29^. It should be noted that the PFS following ICB therapy information was not available for this cohort.

### TMB and NLR harmonization

TMB for different cohorts were determined using two different platforms, namely, whole exome sequencing (the *Shim et al., Ravi et al., and Pradat et al.* cohorts) and targeted tumor-sequencing (other cohorts). We harmonized whole exome sequencing TMB values to MSK-IMPACT panel TMB values. To do this, we first merged all MSK cohorts into one, comprising the *Chowell et al., MSK1, MSK2, Vanguri et al.* cohorts. Then, we normalized cohort-specific TMB distributions using power transformations and the R package *rcompanion*’s function *transformTukey*(). Thirdly, we standardized the distributions to Z-scores by subtracting the transformed distribution mean and dividing by the standard deviation, as previously described ^45^. To ensure the final TMB values held biological relevance, we converted all Z-scored TMB values from whole exome sequencing cohorts back to the level of MSK-measured TMB values. This was achieved using transformation coefficients calculated from the consolidated MSK cohort. It is noteworthy that, for the standardization of the NSCLC-specific *Shim et al.* and *Ravi et al.* cohorts, only NSCLC patients from the merged MSK cohort were utilized for the calculation of transformation coefficients.

Derived NLR (dNLR) was measured for the *Vanguri et al.* cohort. To harmonize it with the NLR in the merged MSK cohort, we employed a strategy similar to that used for TMB standardization. In this case, too, only NSCLC patients from the merged MSK cohort were used for calculating transformation coefficients.

### Patient outcome following immunotherapy

Patient outcome following immunotherapy was assessed by measuring objective response, OS, and PFS for all cohorts described above. Objective response was categorized based on the RECIST v1.1 criteria ^46^, except for the CNS tumors in the *MSK2* cohort, where the Response Assessment in Neuro-Oncology (RANO) criteria were used instead ^47^. The objective response was then dichotomized into responders (CR and PR) and non-responders (SD and PD). PFS was defined as the time from the first infusion of ICB to disease progression or death from any cause. Patients without disease progression were censored at their last disease assessment. OS was defined as the time from the first ICB infusion to death from any cause, and patients who were still alive at the time of review were censored at their last contact.

### Developing multivariable models of ICB response

No data imputation was performed during the development and testing of all models and biomarkers. Patients with incomplete data were excluded from corresponding models. To minimize the impact of extreme values in some features, data truncation was applied. TMB values were truncated at 50 Mut/Mb, blood NLR was truncated at 25, and patient age was truncated at 85 years.

#### Pan-cancer study

We utilized a subset of patients from the *Chowell et al.* cohort who underwent immunotherapy between 2015 and 2017 as our training set (*Chowell training*, n=964). We investigated 20 machine learning classifiers to predict patient ICB response, using the FDA-approved TMB biomarker as a baseline model. Among these models, the decision tree classifier and random forest classifiers directly took the raw feature values as input. For all other classifiers, all feature values (continuous and binary) were standardized by converting them to z-scores (i.e., mean normalized to zero and standard deviation normalized to one) before inputting to the models. We built, tuned, and evaluated all the multivariable machine-learning models using the *scikit-learn* package (v.1.2.1) and *keras* (v2.8.0) in the Python programming language (https://www.python.org/, v.3.9.16). We determined the optimal hyperparameter combination by employing a random search approach with the *RandomizedSearchCV* function to maximize the AUC scores in a five-fold cross-validation of the training dataset data. We determined the total number of different hyperparameter combinations for each model, N, as the minimum of 10,000 and the total number of all possible combinations. The combinations of hyperparameters and the identified optimal combination for each of the 20 machine learning classifiers are listed below:

1. The 16-feature logistic regression classifier using all 16 features measured in the *Chowell et al.* cohort. All combinations: solver = “saga”; penalty = “elasticnet”; class_weight = “balanced”; l1_ratio from 0 to 1 (step size 0.1); max_iter from 100 to 1,000 (step size 100); C from 10^-3^ to 10^3^ (logarithmic step size 1). Optimal combination: solver = “saga”, penalty = “elasticnet”, max_iter = 100, l1_ratio = 0.1, class_weight = “balanced”, C = 0.01.
2. The 6-feature logistic regression classifier using TMB, patient systemic therapy history, blood albumin level, blood NLR, patient age, and cancer type. All combinations: same as above. Optimal combination: solver = “saga”, penalty = “elasticnet”, max_iter = 100, l1_ratio = 1, class_weight = “balanced”, C = 0.1.
3. The 5-feature logistic regression classifier subtracting TMB from the previous model. All combinations: same as above. Optimal combination: solver = “saga”, penalty = “elasticnet”, max_iter = 100, l1_ratio = 0.4, class_weight = “balanced”, C = 0.01.
4. The 16-feature random forest classifier with hyperparameters reported by *Chowell et al.* ^22^. Optimal combination: n_estimators=1,000, max_depth=8, min_samples_leaf=20, min_samples_split=2.
5. The 16-feature random forest classifier re-trained using protocol in this study. All combinations: n_estimators from 200 to 2,000 (step size 200); max_features from 0.1 to 0.9 (step size 0.1); max_depth from 3 to 10 (step size 1); min_samples_leaf from 2 to 30 (step size 2); min_samples_split from 2 to 30 (step size 2). Optimal combination: n_estimators = 400, max_features = 0.1, max_depth = 9, min_samples_leaf = 2, min_samples_split = 8.
6. The 6-feature random forest classifier trained using protocol by *Chowell et al.* ^22^. All combinations: n_estimators from 100 to 1,000 (step size 100); max_depth from 2 to 20 (step size 2); min_samples_leaf from 2 to 20 (step size 2); min_samples_split from 2 to 20 (step size 2). Optimal combination: n_estimators = 900, max_depth = 8, min_samples_leaf = 8, min_samples_split = 20.
7. The decision tree classifier. All combinations: splitter = “best” or “random”; max_features from 0.1 to 0.9 (step size 0.1); max_depth from 3 to 10 (step size 1); min_samples_leaf from 2 to 30 (step size 2); min_samples_split from 2 to 30 (step size 2); ccp_alpha = 0 or 0.5 or 1 or 10 or 100. Optimal combination: splitter = random, max_features = 0.7, max_depth = 7, min_samples_leaf = 8, min_samples_split = 2, ccp_alpha = 0.
8. The GBoost classifier. All combinations: learning_rate = 0.01 or 0.03 or 0.05 or 0.1 or 0.3 or 0.5; n_estimators from 200 to 2,000 (step size 200); min_samples_split from 2 to 30 (step size 2); min_samples_leaf from 2 to 30 (step size 2); max_depth from 3 to 10 (step size 1); max_features from 0.1 to 0.9 (step size 0.1). Optimal combination: learning_rate = 0.03, n_estimators = 200, min_samples_split = 12, min_samples_leaf = 4, max_depth = 6, max_features = 0.1.
9. The AdaBoost classifier. All combinations: n_estimators from 200 to 2,000 (step size 200); learning_rate = 0.01 or 0.05 or 0.03 or 0.1 or 0.3 or 0.5 or 1; algorithm = “SAMME” or “SAMME.R”. Optimal combination: n_estimators = 1000, learning_rate = 0.3, algorithm = SAMME.
10. The HGBoost classifier. All combinations: learning_rate = 0.01 or 0.03 or 0.05 or or 0.3 or 0.5; max_iter from 200 to 2,000 (step size 200); min_samples_leaf from 2 to 30 (step size 2); max_depth from 3 to 10 (step size 1); l2_regularization = 0 or from 10^-4^ to 10^2^ (logarithmic step size 1). Optimal combination: learning_rate = 0.03, max_iter = 600, min_samples_leaf = 16, max_depth = 10, l2_regularization = 100.
11. The XGBoost classifier. All combinations: min_child_weight = 1 or from 2 to 30 (step size 2); max_depth from 3 to 10 (step size 1); n_estimators = 100 or from 200 to 1,000 (step size 200); learning_rate = 0.01 or 0.03 or 0.05 or 0.1 or 0.3 or 0.5; colsample_bytree = 0.5 or 0.8 or 1; colsample_bynode from 0.2 to 1 (step size 0.2); colsample_bylevel from 0.2 to 1 (step size 0.2). Optimal combination: min_child_weight = 6, max_depth = 7, n_estimators = 400, learning_rate = 0.01, colsample_bytree = 0.8, colsample_bynode = 0.2, colsample_bylevel = 1.
12. The LightGBM classifier. All combinations: learning_rate = 0.001 or 0.003 or 0.005 or 0.01 or 0.03 or 0.05 or 0.1 or 0.3; max_depth from 3 to 10 (step size 1); n_estimators from 200 to 2,000 (step size 200); num_leaves from 10 to 100 (step size 10); colsample_bytree from to 1 (step size 0.2); min_data_in_leaf from 2 to 30 (step size 2). Optimal combination: learning_rate = 0.03, max_depth = 3, n_estimators = 200, num_leaves = 30, colsample_bytree = 0.8, min_data_in_leaf = 30.
13. The support vector machine classifier. All combinations: C from 10^-5^ to 10^3^ (logarithmic step size 0.5); gamma = “scale” or “auto” or from 10^-4^ to 10^2^ (logarithmic step size 0.5); kernel = “rbf”; max_iter = -1 or100 or1000; tol from 10^-5^ to 10^-1^ (logarithmic step size 0.5); class_weight = None or “balanced”. Optimal combination: C = 10^2.5^, gamma = 10^-3.5^, kernel = “rbf”, max_iter = 1000, tol = 0.001, class_weight = None.
14. The k-nearest neighbors classifier. All combinations: n_neighbors from 2 to 60 (step size 2); weights = “uniform” or “distance”; algorithm = “auto” or “ball_tree” or “kd_tree” or “brute”; leaf_size from 2 to 30 (step size 2); p from 1 to 10 (step size 1). Optimal combination: n_neighbors = 58, weights = “distance”, algorithm = “brute”, leaf_size = 20, p = 1.
15. The Keras sequential classifier. All combinations: layers = [16] or [32, 16] or [64, 32, 16]; activation = “sigmoid” or “relu”; batch_size = 32 or 64 or 128 or 256; epochs = 10 or 50 or 100. Optimal combination: layers = [64, 32, 16], activation = “sigmoid”, batch_size = 32, epochs = 100.
16. The multilayer perceptron classifier (1 layer) . All combinations: solver = “sgd” or “lbfgs” or “adam”; learning_rate = “constant” or “invscaling” or “adaptive”; max_iter = 100 or 200 or 500 or 1,000; hidden_layer_sizes from 2 to 40 (step size 1) in one hidden layer; activation = “logistic” or “tanh” or “relu” or “identity”; alpha from 10^-6^ to 10^-1^ (logarithmic step size 1); early_stopping = False or True. Optimal combination: solver = “adam”, learning_rate = “adaptive”, max_iter = 200, hidden_layer_sizes = (19,), activation = “tanh”, alpha = 10^-2^, early_stopping = False.
17. The multilayer perceptron classifier (2 layers). All combinations: max_iter = 100 or 200 or 500 or 1,000; hidden_layer_sizes from 2 to 20 (step size 1) in two hidden layers; activation = “logistic” or “tanh” or “relu” or “identity”; alpha from 10^-6^ to 10^-1^ (logarithmic step size 1); early_stopping = False or True. Optimal combination: max_iter = 100, hidden_layer_sizes = (19, 19), activation = “tanh”, alpha = 10^-5^, early_stopping = False.
18. The multilayer perceptron classifier (3 layers). All combinations: hidden_layer_sizes from 2 to 20 (step size 1) in three hidden layers; activation = “logistic” or “tanh” or “relu” or “identity”; alpha from 10^-6^ to 10^-1^ (logarithmic step size 1). Optimal combination: hidden_layer_sizes = (6, 5, 6), activation = “relu”, alpha = 10^-1^.
19. The multilayer perceptron classifier (4 layers). All combinations: same as above. Optimal combination: hidden_layer_sizes = (3, 17, 2, 4), activation = “tanh”, alpha = 10^-3^.
20. The Gaussian process classifier. All combinations: kernel = None or 1.0 * kernels.RBF(1.0) or 0.1 * kernels.RBF(0.1) or 10 * kernels.RBF(10); optimizer = “fmin_l_bfgs_b” or None; max_iter_predict = 100 or 500 or 1,000; n_restarts_optimizer from 0 to 30 (step size 5). Optimal combination: kernel = 10 * RBF(length_scale=10), optimizer = None, max_iter_predict = 100, n_restarts_optimizer = 0.

After hyperparameter tuning, it was observed that the logistic regression model with 6 features had a LASSO penalty proportion of 100%, making it a logistic LASSO regression (LLR) model. For ease of reference, we referred to this model as LLR6 throughout the paper. The hyperparameters that were optimal for LLR6 were used to train the NSCLC-specific model. The regression coefficients, which include the intercept, from the pan-cancer LLR6 model were obtained by averaging the corresponding values obtained from the 10,000 training iterations described earlier. No further adaptation was done on the test data. The LLR6 score, calculated using this model, was referred to as LORIS. The formulation for pan-cancer LORIS is explicitly given as follows:

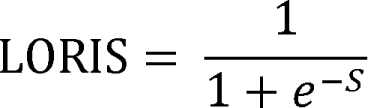

*S = 0.0371· min (MB, 50) - 0.8775 · Chemo + 0.5382 · Alb min - 0.033 · min(NLR, 25) + 0.0049 · min(Age, 85) - 0.3323 · Bladder - 0.3323 · Breast - 0.102 · Colorectal - 0.0079 · Endometrial + 0.55 · Esophageal + 0.2306 · Gastric + 0.0678 · Head &Neck - 0.1189 · Hepatobiliary - 0.0086 · Melanoma + 0.1255 · Mesothelioma + 0.0008 · NSCLC - 0.052 · Ovarian - 1.1169 · Pancreatic + 0.5451 · Renal + 0.0542 · Sarcoma - 0.0033 · SCLC - 2.0886*

#### NSCLC study

The NSCLC-specific study aimed to replicate the pan-cancer study, with the difference that only NSCLC patients were used for training and evaluating the models. We used the *Chowell et al.* cohort as training data, which included 324 NSCLC patients.

As previously mentioned, we used the optimal hyperparameters obtained from the pan-cancer study to train the NSCLC-specific LLR6 model. We followed a similar approach to the pan-cancer modeling approach, calculating coefficients and intercepts for NSCLC-specific LLR6 based on the average values of 10,000 training iterations, with random 80% training data used for each iteration. These values were then applied directly to the entire test dataset. The formula for NSCLC-specific LLR6 in calculating LORIS is as follows:

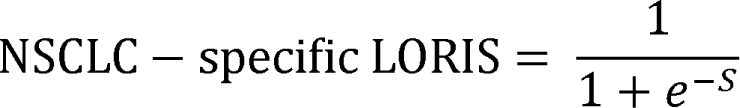

*S = 0.0353· min (MB, 50) + 0.0111 · P>L1 - 0.375 · Chemo + 0.2924 · Alb min - 0.0103· min(NLR, 25) + 0 · min(Age, 85) - 1.5593*

### Model performance evaluation

To evaluate the performance of the models, we employed 2,000-repeated 5-fold cross-validation on the training data. During each cross-validation fold, 80% of the training data was used for model training, and the remaining 20% was used for evaluation. We utilized several metrics such as accuracy, F1 score, confusion matrix, and odds ratio (of response) to quantify the predictive power of the models. To determine the optimal threshold for the predicted response probabilities computed by the model, we maximized the Youden’s index, defined as “sensitivity + specificity – 1”. We evaluated the performance of each model using the geometric mean of four metrics: AUC, AUPRC, accuracy, and F1 score. As overfitting is a common problem in supervised models, we calculated the difference between the performance scores on the training data and cross-validation data for each cross-validation fold to estimate the extent of overfitting for each model.

### Statistical analyses

We conducted various statistical analyses using Python (v.3.9.5) and R (v.4.1.1) to investigate the relationships between different variables and ICB response. Spearman’s rank test from the *scipy* package (v.1.10.1) was used to calculate correlation coefficients and raw p values between features measured on a continuous scale, which were then adjusted for Bonferroni correction. To compare the distributions of response probability generated by different models (e.g., LLR6, RF6, and TMB) between responders and non-responders, we used the Mann-Whitney U test.

Survival analysis was performed using the R packages *survminer* v.0.4.9 and *survival* v.3.3.1, and we calculated HR and P values with univariable Cox proportional hazard regression using the *coxph*() function 19. To compare differences in half-year, 1-year, 2-year, 3-year, 4-year, and 5-year survival probability between high and low LORIS or TMB groups in individual cancer types, we used paired Wilcoxon test. Multivariable analysis was performed with Cox proportional hazard regression in individual cancer types using the *coxph*() function 19, with adjustment for cancer type, age, drug class of ICB, and year of ICB start.

For variables such as LORIS, PD-L1 TPS, and TMB, we calculated the average values and 95% confidence intervals using 1,000-replicate bootstrapping of the data to determine their relationship with ICB objective response probability. As the features were z-score normalized before being put into the model, the feature importance for the logistic regression model was directly shown as the absolute values of the corresponding coefficients in the model.

All statistical tests were two-sided unless otherwise stated.

## Code availability

All codes that are necessary to reproduce all the results in the paper are implemented in Python and R and are publicly available in GitHub: https://github.com/rootchang/ICBpredictor.

## Data availability

All deidentified data required to replicate our analyses will be made publicly available as a supplementary table upon acceptance of the manuscript.

## Acknowledgements

This work utilized the computational resources of the NIH HPC Biowulf cluster (http://hpc.nih.gov).

## Author contributions

T-G.C. and E.R. conceived and designed the study. T-G.C., Y.C. and S.R.D. developed the machine learning model. T-G.C., Y.C., S.R.D., S-H.L., C.V., S-K.Y., D.C., L.G.T.M, and E.R. acquired, analyzed or interpreted the data. All authors critically revised the manuscript for important intellectual content. E.R. and L.G.T.M. supervised the study.

## References

1. Topalian, S.L., Taube, J.M., Anders, R.A. & Pardoll, D.M. Mechanism-driven biomarkers to guide immune checkpoint blockade in cancer therapy. Nature Reviews Cancer 16, 275–287 (2016).

2. Morad, G., Helmink, B.A., Sharma, P. & Wargo, J.A. Hallmarks of response, resistance, and toxicity to immune checkpoint blockade. Cell 184, 5309–5337 (2021).

3. Nishino, M., Ramaiya, N.H., Hatabu, H. & Hodi, F.S. Monitoring immune-checkpoint blockade: response evaluation and biomarker development. Nature Reviews Clinical Oncology 14, 655–668 (2017).

4. Goodman, A.M. et al. Tumor Mutational Burden as an Independent Predictor of Response to Immunotherapy in Diverse Cancers. Molecular Cancer Therapeutics 16, 2598–2608 (2017).

5. Samstein, R.M. et al. Tumor mutational load predicts survival after immunotherapy across multiple cancer types. Nature Genetics 51, 202-+ (2019).

6. McGrail, D.J. et al. High tumor mutation burden fails to predict immune checkpoint blockade response across all cancer types. Annals of Oncology 32, 661–672 (2021).

7. Topalian, S.L. et al. Safety, Activity, and Immune Correlates of Anti-PD-1 Antibody in Cancer. New England Journal of Medicine 366, 2443–2454 (2012).

8. Zhao, P.F., Li, L., Jiang, X.Y. & Li, Q. Mismatch repair deficiency/microsatellite instability-high as a predictor for anti-PD-1/PD-L1 immunotherapy efficacy. Journal of Hematology & Oncology 12(2019).

9. Mandal, R. et al. CANCER Genetic diversity of tumors with mismatch repair deficiency influences anti-PD-1 immunotherapy response. Science 364, 485-+ (2019).

10. Le, D.T. et al. Mismatch repair deficiency predicts response of solid tumors to PD-1 blockade. Science 357, 409–413 (2017).

11. Chowell, D. et al. Evolutionary divergence of HLA class I genotype impacts efficacy of cancer immunotherapy. Nature Medicine 25, 1715-+ (2019).

12. Chowell, D. et al. Patient HLA class I genotype influences cancer response to checkpoint blockade immunotherapy. Science 359, 582-+ (2018).

13. Davoli, T., Uno, H., Wooten, E.C. & Elledge, S.J. Tumor aneuploidy correlates with markers of immune evasion and with reduced response to immunotherapy. Science 355(2017).

14. Chang, T., Cao, Y., Shulman, E.D., Schäffer, A.A. & Ruppin, E. Fraction of copy-number alterations significantly predicts survival following immunotherapy in a few cancers. bioRxiv, 2022.12. 28.522101 (2022).

15. Ren, F.P., Zhao, T., Liu, B. & Pan, L. Neutrophil-lymphocyte ratio (NLR) predicted prognosis for advanced non-small-cell lung cancer (NSCLC) patients who received immune checkpoint blockade (ICB). Oncotargets and Therapy 12, 4235–4244 (2019).

16. Valero, C. et al. Pretreatment neutrophil-to-lymphocyte ratio and mutational burden as biomarkers of tumor response to immune checkpoint inhibitors. Nature Communications 12(2021).

17. Yoo, S.K., Chowell, D., Valero, C., Morris, L.G.T. & Chan, T.A. Pre-treatment serum albumin and mutational burden as biomarkers of response to immune checkpoint blockade. Npj Precision Oncology 6(2022).

18. Wang, Z.M. et al. Paradoxical effects of obesity on T cell function during tumor progression and PD-1 checkpoint blockade. Nature Medicine 25, 141-+ (2019).

19. Conforti, F. et al. Cancer immunotherapy efficacy and patients’ sex: a systematic review and meta-analysis. Lancet Oncology 19, 737–746 (2018).

20. Kugel, C.H. et al. Age Correlates with Response to Anti-PD1, Reflecting Age-Related Differences in Intratumoral Effector and Regulatory T-Cell Populations. Clinical Cancer Research 24, 5347–5356 (2018).

21. Litchfield, K. et al. Meta-analysis of tumor- and T cell-intrinsic mechanisms of sensitization to checkpoint inhibition. Cell 184, 596-+ (2021).

22. Chowell, D. et al. Improved prediction of immune checkpoint blockade efficacy across multiple cancer types. Nature Biotechnology 40, 499-+ (2022).

23. Gromeier, M. et al. Very low mutation burden is a feature of inflamed recurrent glioblastomas responsive to cancer immunotherapy. Nature Communications 12(2021).

24. Diggs, L.P. & Hsueh, E.C. Utility of PD-L1 immunohistochemistry assays for predicting PD-1/PD-L1 inhibitor response. Biomarker Research 5(2017).

25. Shim, J.H. et al. HLA-corrected tumor mutation burden and homologous recombination de fi ciency for the prediction of response to PD -(L)1 blockade in advanced non -small - cell lung cancer patients. Annals of Oncology 31, 902–911 (2020).

26. Kato, S. et al. Real-world data from a molecular tumor board demonstrates improved outcomes with a precision N-of-One strategy. Nature Communications 11(2020).

27. Vanguri, R.S. et al. Multimodal integration of radiology, pathology and genomics for prediction of response to PD-(L)1 blockade in patients with non-small cell lung cancer. Nature Cancer 3, 1151-+ (2022).

28. Ravi, A. et al. Genomic and transcriptomic analysis of checkpoint blockade response in advanced non-small cell lung cancer. Nature Genetics (2023).

29. Pradat, Y. et al. Integrative Pan-Cancer Genomic and Transcriptomic Analyses of Refractory Metastatic Cancer. Cancer Discovery 13, 1116–1143 (2023).

30. Eisenhauer, E.A. et al. New response evaluation criteria in solid tumours: Revised RECIST guideline (version 1.1). European Journal of Cancer 45, 228–247 (2009).

31. Cho, M.S. et al. Platelets Increase the Expression of PD-L1 in Ovarian Cancer. Cancers 14(2022).

32. Rudin, C. Stop explaining black box machine learning models for high stakes decisions and use interpretable models instead. Nature Machine Intelligence 1, 206–215 (2019).

33. Petch, J., Di, S. & Nelson, W. Opening the Black Box: The Promise and Limitations of Explainable Machine Learning in Cardiology. Can J Cardiol 38, 204–213 (2022).

34. Watson, D.S. et al. Clinical applications of machine learning algorithms: beyond the black box. Bmj-British Medical Journal 364(2019).

35. Sambi, M., Bagheri, L. & Szewczuk, M.R. Current Challenges in Cancer Immunotherapy: Multimodal Approaches to Improve Efficacy and Patient Response Rates. Journal of Oncology 2019(2019).

36. He, Y.Y. et al. Genomic and transcriptional alterations in first-line chemotherapy exert a potentially unfavorable influence on subsequent immunotherapy in NSCLC. Theranostics 11, 7092–7109 (2021).

37. Haas, L. et al. Acquired resistance to anti-MAPK targeted therapy confers an immune-evasive tumor microenvironment and cross-resistance to immunotherapy in melanoma. Nature Cancer 2, 693-+ (2021).

38. Razavian, N. et al. Population-Level Prediction of Type 2 Diabetes From Claims Data and Analysis of Risk Factors. Big Data 3, 277–287 (2015).

39. Bareche, Y. et al. Leveraging big data of immune checkpoint blockade response identifies novel potential targets. Annals of Oncology 33, 1304–1317 (2022).

40. Cercek, A. et al. PD-1 Blockade in Mismatch Repair-Deficient, Locally Advanced Rectal Cancer. New England Journal of Medicine 386, 2363–2376 (2022).

41. Negrao, M.V. et al. PD-L1 Expression, Tumor Mutational Burden, and Cancer Gene Mutations Are Stronger Predictors of Benefit from Immune Checkpoint Blockade than HLA Class I Genotype in Non-Small Cell Lung Cancer. Journal of Thoracic Oncology 14, 1021–1031 (2019).

42. Auslander, N. et al. Robust prediction of response to immune checkpoint blockade therapy in metastatic melanoma (vol 24, pg 1545, 2018). Nature Medicine 24, 1942–1942 (2018).

43. Jiang, P. et al. Signatures of T cell dysfunction and exclusion predict cancer immunotherapy response. Nature Medicine 24, 1550-+ (2018).

44. Liu, D. et al. Integrative molecular and clinical modeling of clinical outcomes to PD1 blockade in patients with metastatic melanoma (vol 25, pg 1915, 2019). Nature Medicine 26, 1147–1147 (2020).

45. Vokes, N.I., et al. Harmonization of Tumor Mutational Burden Quantification and Association With Response to Immune Checkpoint Blockade in Non-Small-Cell Lung Cancer. Jco Precision Oncology 3(2019).

46. Schwartz, L.H. et al. RECIST 1.1-Update and clarification: From the RECIST committee. European Journal of Cancer 62, 132–137 (2016).

47. Wen, P.Y. et al. Updated Response Assessment Criteria for High-Grade Gliomas: Response Assessment in Neuro-Oncology Working Group. Journal of Clinical Oncology 28, 1963–1972 (2010).

