## Supplementary information for "Robust prediction of patient outcomes with immune checkpoint blockade therapy for cancer using common clinical, pathologic, and genomic features"

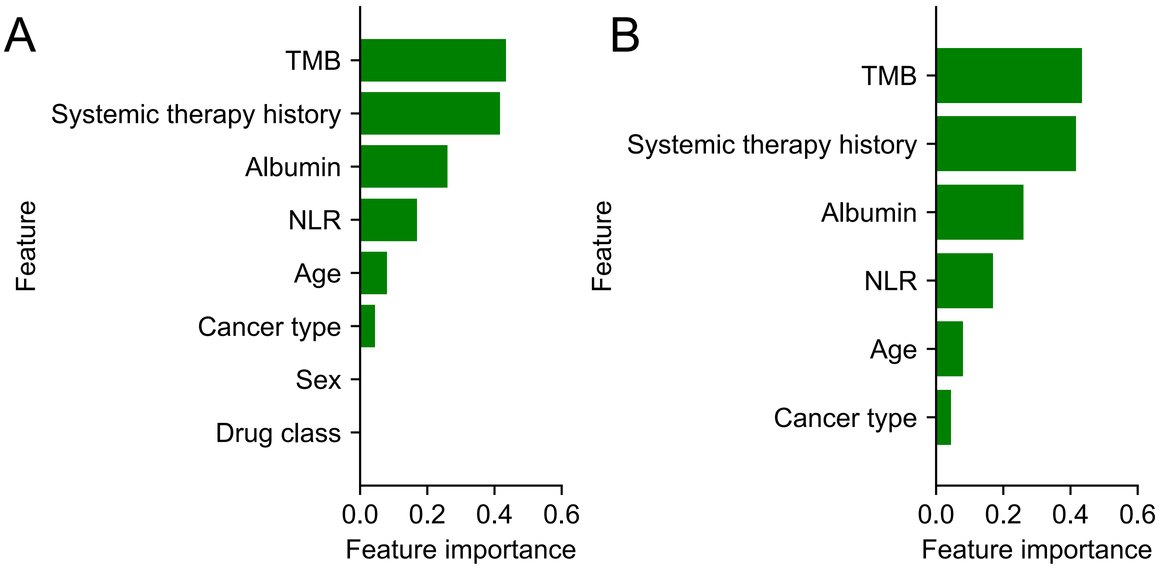


**Supplementary Figure 1. Feature selection and feature importance by the logistic LASSO regression model.**

**A**. Feature importance of from the 8-feature logistic regression classifier using features commonly measured across most patients.

**B**. Feature importance of the 6-feature logistic regression classifier LLR6. The feature importance is calculated as the absolute values of the corresponding coefficients in the logistic regression models. Importance for cancer type is calculated as the average importance of individual cancer types.

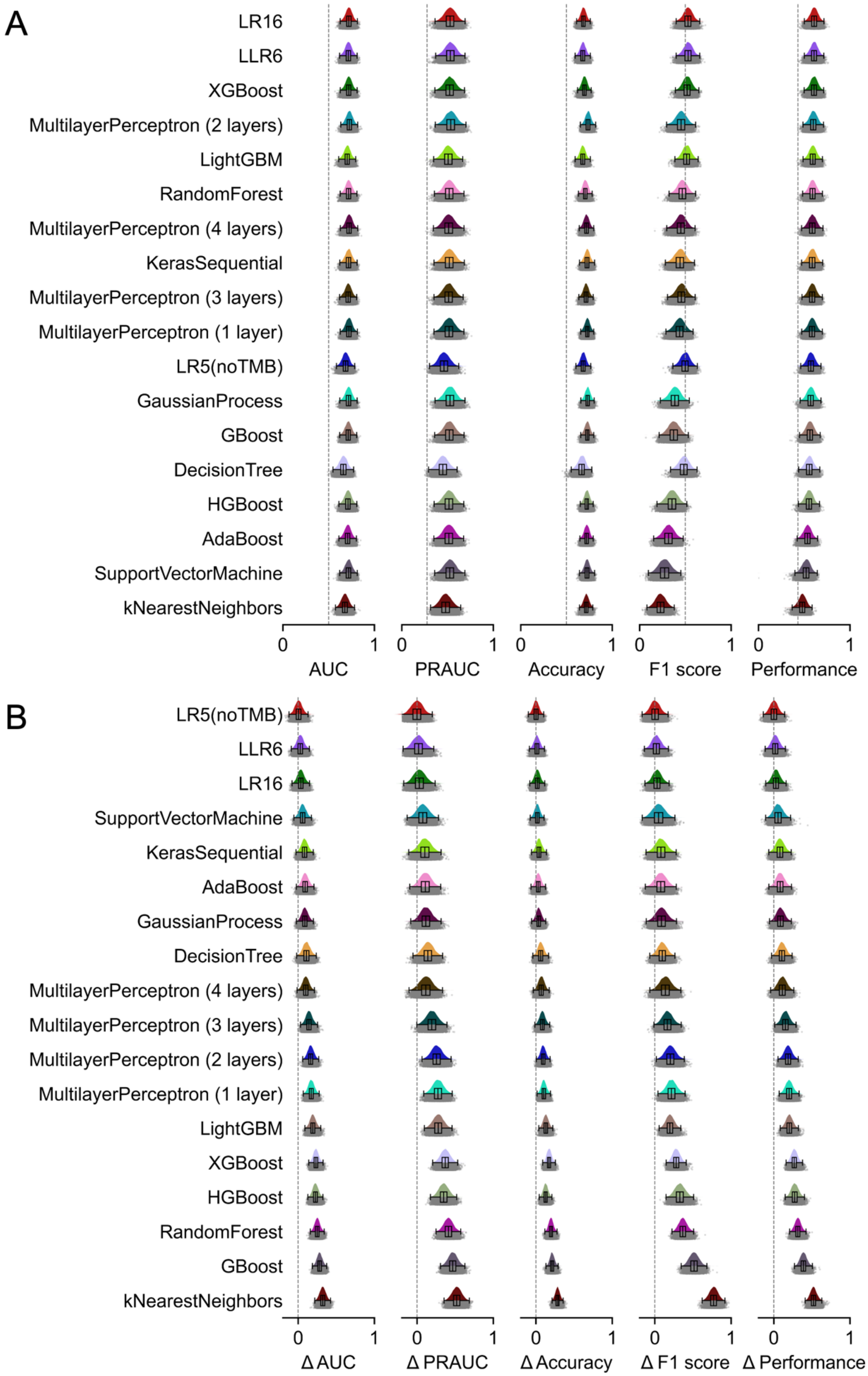
A
LR16
LLR6
XGBoost
MultilayerPerceptron (2 layers)
LightGBM
RandomForest
MultilayerPerceptron (4 layers)
KerasSequential
MultilayerPerceptron (3 layers)
MultilayerPerceptron (1 layer)
LR5(noTMB)
GaussianProcess
GBoost
DecisionTree
HGBoost
AdaBoost
SupportVectorMachine
kNearestNeighbors
0 1 0 1 0 1 0 1 0 1
AUC
PRAUC
Accuracy
F1 score
Performance
B
LR5(noTMB)
LLR6
LR16
SupportVectorMachine
KerasSequential
AdaBoost
GaussianProcess
DecisionTree
MultilayerPerceptron (4 layers)
MultilayerPerceptron (3 layers)
MultilayerPerceptron (2 layers)
MultilayerPerceptron (1 layer)
LightGBM
XGBoost
HGBoost
RandomForest
GBoost
kNearestNeighbors
0 1 0 1 0 1 0 1 0 1
Δ AUC
Δ PRAUC
Δ Accuracy
Δ F1 score
Δ Performance

**Supplementary Figure 2. Comparison of predictive power of pan-cancer models for ICB response prediction.**

**A**. Model predictive power on the validation data. Model *performance* is defined as the geometric mean of four metrics, i.e., AUC, AUPRC, accuracy, and F1 score.

**B**. Difference of model predictive power between the training and the validation data Various performance metrics were calculated with 2000-repeated 5-fold cross-validation.

All machine learning models take all 16 measured features from the *Chowell et al.* cohort as input, with the exception of the LLR6 and LR5 models, which utilize 6 and 5 features, respectively.

In the figure, the grey dots represent individual model scores from each fold during cross-validation. The colored histograms illustrate the distributions of these model scores across all folds of the cross-validation process. Regarding the box plots, the upper and lower boundaries signify the first and third quartiles, correspondingly, the central line denotes the median, and the whiskers stretch to the most distant data points not classified as outliers (within 1.5 times the interquartile range).

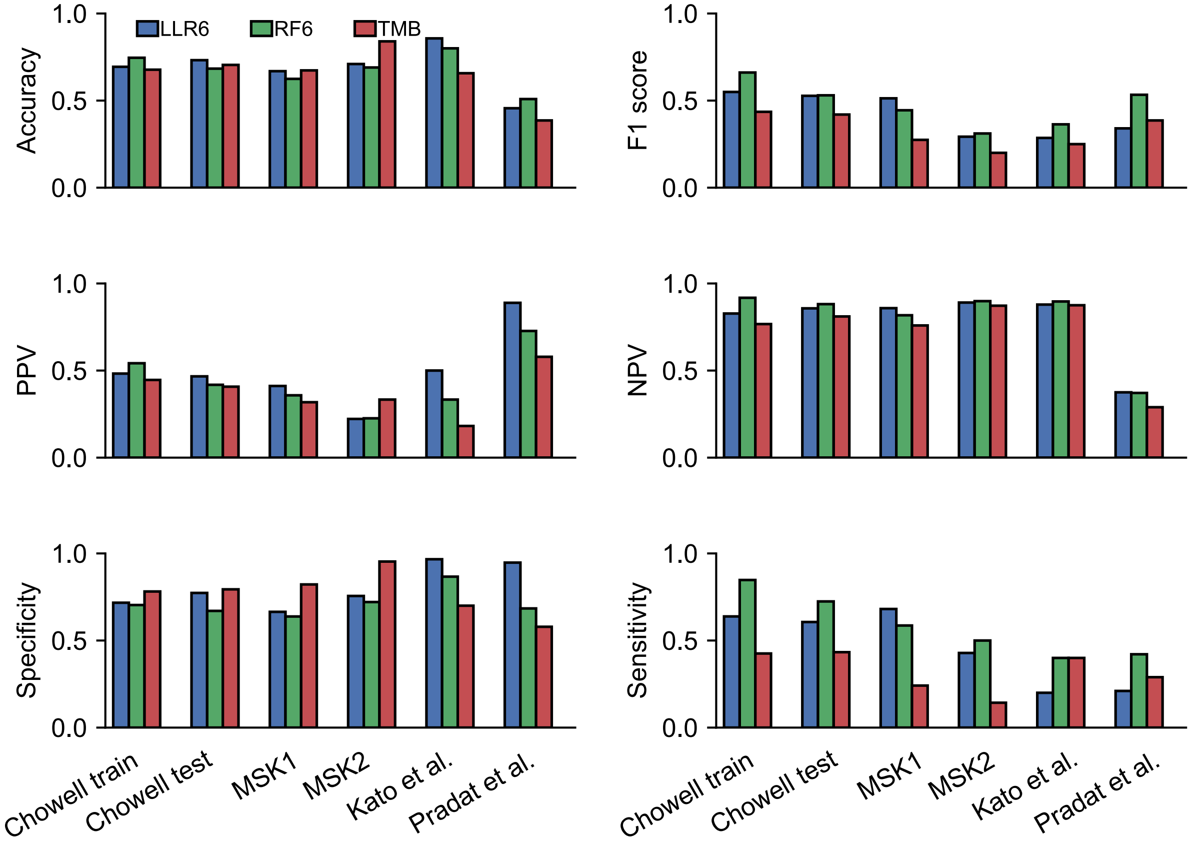


**Supplementary Figure 3. Comparison of model predictive power on the training set and across multiple unseen test sets.**

Multiple metrics were used to compare predictive power of LLR6, RF6, and the TMB biomarker, including accuracy, F1 score, positive predictive value (PPV), negative predictive value (NPV), specificity, and sensitivity.

The optimal thresholds of 0.5 and 0.27 that differentiate responders from non-responders were determined on the training data for LLR6 and RF6 models using the Youden's index respectively. As for the TMB model, the threshold of 10 Mut/Mb was utilized.

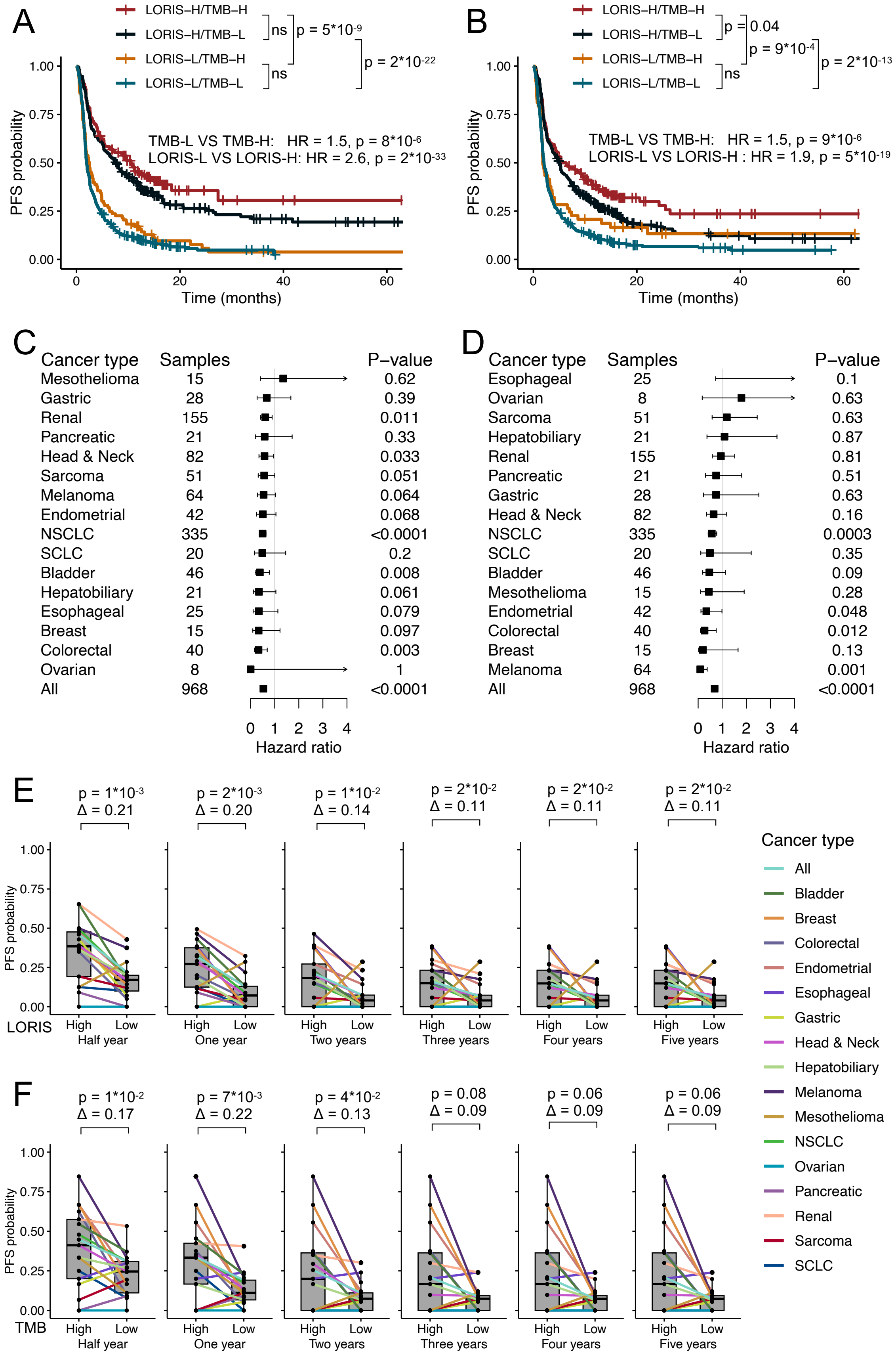
A
LORIS–H/TMB–H
LORIS–H/TMB–L
LORIS–L/TMB–H
LORIS–L/TMB–L
ns
p = 5*10-9
p = 2*10-22
ns
TMB-L VS TMB-H: HR = 1.5, p = 8*10-6
LORIS-L VS LORIS-H: HR = 2.6, p = 2*10-33
PFS probability
Time (months)
B
LORIS–H/TMB–H
LORIS–H/TMB–L
LORIS–L/TMB–H
LORIS–L/TMB–L
p = 0.04
p = 9*10-4
p = 2*10-13
ns
TMB-L VS TMB-H: HR = 1.5, p = 9*10-6
LORIS-L VS LORIS-H : HR = 1.9, p = 5*10-19
PFS probability
Time (months)
C
Cancer type Samples P–value
Mesothelioma 15 0.62
Gastric 28 0.39
Renal 155 0.011
Pancreatic 21 0.33
Head & Neck 82 0.033
Sarcoma 51 0.051
Melanoma 64 0.064
Endometrial 42 0.068
NSCLC 335 <0.0001
SCLC 20 0.2
Bladder 46 0.008
Hepatobiliary 21 0.061
Esophageal 25 0.079
Breast 15 0.097
Colorectal 40 0.003
Ovarian 8 1
All 968 <0.0001
Hazard ratio
D
Cancer type Samples P–value
Esophageal 25 0.1
Ovarian 8 0.63
Sarcoma 51 0.63
Hepatobiliary 21 0.87
Renal 155 0.81
Pancreatic 21 0.51
Gastric 28 0.63
Head & Neck 82 0.16
NSCLC 335 0.0003
SCLC 20 0.35
Bladder 46 0.09
Mesothelioma 15 0.28
Endometrial 42 0.048
Colorectal 40 0.012
Breast 15 0.13
Melanoma 64 0.001
All 968 <0.0001
Hazard ratio
E
p = 1*10-3 Δ = 0.21
p = 2*10-3 Δ = 0.20
p = 1*10-2 Δ = 0.14
p = 2*10-2 Δ = 0.11
p = 2*10-2 Δ = 0.11
p = 2*10-2 Δ = 0.11
PFS probability
LORIS High Low
Half year
One year
Two years
Three years
Four years
Five years
Cancer type
All
Bladder
Breast
Colorectal
Endometrial
Esophageal
Gastric
Head & Neck
Hepatobiliary
Melanoma
Mesothelioma
NSCLC
Ovarian
Pancreatic
Renal
Sarcoma
SCLC
F
p = 1*10-2 Δ = 0.17
p = 7*10-3 Δ = 0.22
p = 4*10-2 Δ = 0.13
p = 0.08 Δ = 0.09
p = 0.06 Δ = 0.09
p = 0.06 Δ = 0.09
PFS probability
TMB High Low
Half year
One year
Two years
Three years
Four years
Five years

**Supplementary Figure 4. LORIS predicts PFS following immunotherapy for both pan-cancer and individual cancer types.**

**A.** Kaplan–Meier analysis of pan-cancer PFS with TMB binned at 10 Mut / Mb and LORIS binned at 0.5. P values next to the legend indicate pairwise comparisons. H, high; L, low.

**B**. Same as panel A, but TMB is binned at the highest 20th percentile, and LORIS is binned at the 50th percentile for each cancer type.

**C-D.** Forest plot of PFS within each cancer type using LORIS (binned at the 50th percentile; **C**) or TMB (binned at the highest 20th percentile; **D**) in a multivariable model with adjustment for cancer type, age, drug class of ICB and year of ICB start. Squares positioned at midpoints symbolize point estimates of HRs, and the accompanying bars indicate 95% confidence intervals.

**E-F.** Comparison of half-year, 1-year, 2-year, 3-year, 4-year, and 5-year PFS stratified by cancer type for high versus low LORIS (50th percentile; **E**) and high versus low TMB (highest 20th percentile; **F**). Survival probability differences and paired Wilcoxon-test p-values are displayed in panels E and F. The upper and lower boundaries signify the first and third quartiles, correspondingly, the central line denotes the median, and the whiskers stretch to the most distant data points not classified as outliers (within 1.5 times the interquartile range).

Data are from combined *Chowell test* (n = 515) and *MSK1* (n = 453) sets.

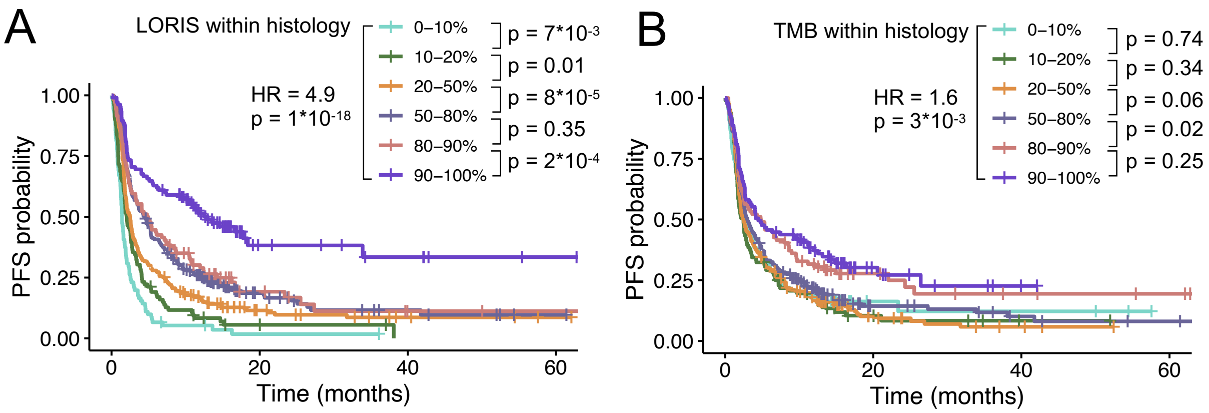


**Supplementary Figure 5. Near-monotonic relationship between LORIS and pan-cancer PFS following immunotherapy.**

Kaplan–Meier analysis of LORIS (**A**) or TMB (**B**) binned at the different percentiles in each cancer type. P values next to the legend indicate pairwise comparisons; hazard ratio (HR) is shown for the lowest-percentile groups (0-10%) and the highest-percentile groups (90-100%). Data are from combined *Chowell test* (n = 515) and *MSK1* (n = 453) sets.

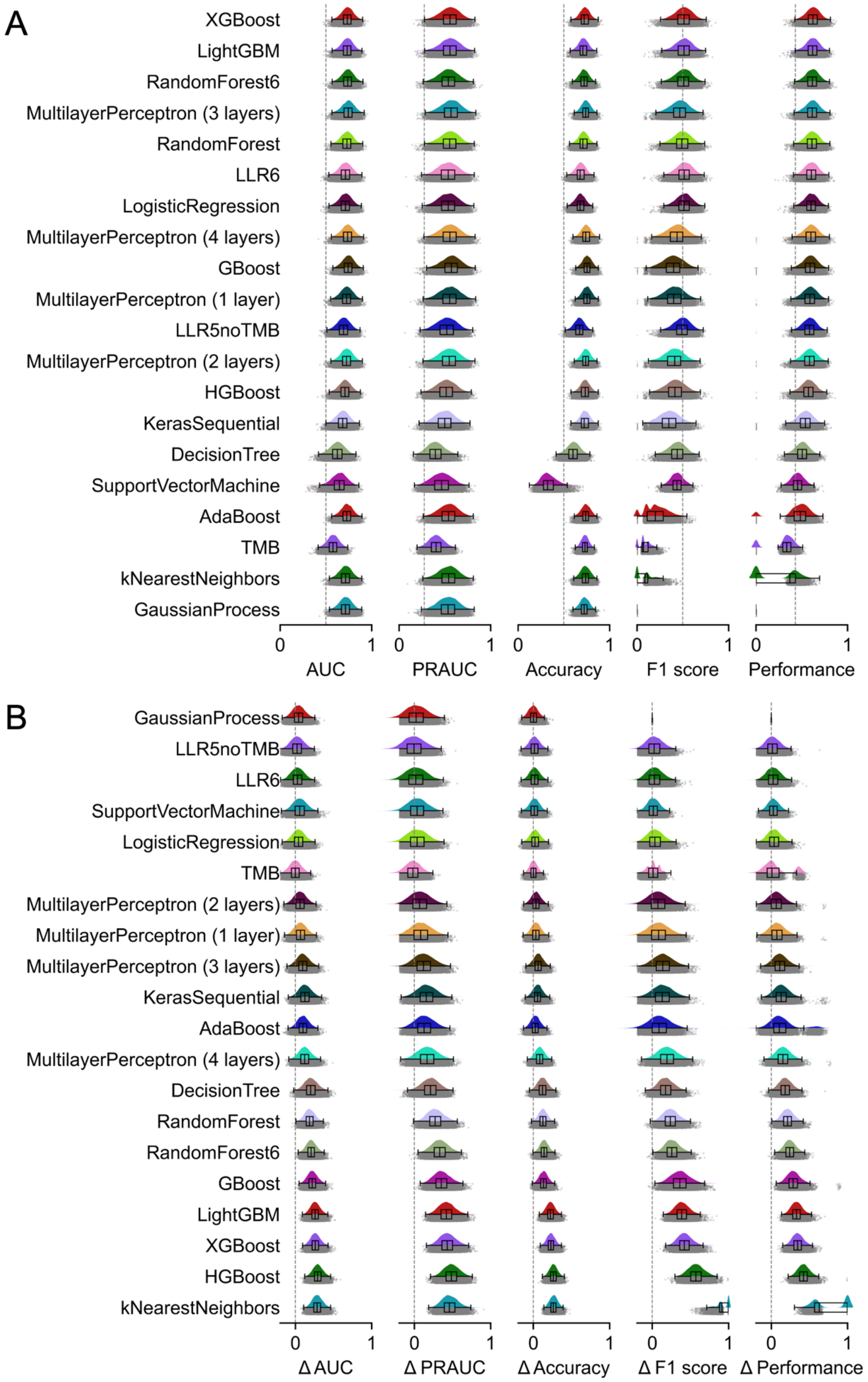
A
XGBoost
LightGBM
RandomForest6
MultilayerPerceptron (3 layers)
RandomForest
LLR6
LogisticRegression
MultilayerPerceptron (4 layers)
GBoost
MultilayerPerceptron (1 layer)
LLR5noTMB
MultilayerPerceptron (2 layers)
HGBoost
KerasSequential
DecisionTree
SupportVectorMachine
AdaBoost
TMB
kNearestNeighbors
GaussianProcess
0 1 0 1 0 1 0 1 0 1
AUC
PRAUC
Accuracy
F1 score
Performance
B
GaussianProcess
LLR5noTMB
LLR6
SupportVectorMachine
LogisticRegression
TMB
MultilayerPerceptron (2 layers)
MultilayerPerceptron (1 layer)
MultilayerPerceptron (3 layers)
KerasSequential
AdaBoost
MultilayerPerceptron (4 layers)
DecisionTree
RandomForest
RandomForest6
GBoost
LightGBM
XGBoost
HGBoost
kNearestNeighbors
0 1 0 1 0 1 0 1 0 1
Δ AUC
Δ PRAUC
Δ Accuracy
Δ F1 score
Δ Performance

**Supplementary Figure 6. Evaluation and comparison of predictive power of NSCLC-specific models.**

**A**. Model predictive power on the validation data. Model *performance* is defined as the geometric mean of four metrics, i.e., AUC, AUPRC, accuracy, and F1 score.

**B**. Difference of model predictive power between the training and the validation data. Various performance metrics were calculated with 2,000-repeated 5-fold cross-validation.

All machine learning models are trained on the *Chowell et al.* cohort using the NSCLC patients and take all 16 measured features as input except the LLR6 and RandomForest6 models, which use 6 features.

In the figure, the grey dots represent individual model scores from each fold during cross-validation. The colored histograms illustrate the distributions of these model scores across all folds of the cross-validation process. Regarding the box plots, the lower and upper boundaries mark the first and third quartiles, respectively, while the central line signifies the median value. The whiskers extend to the furthest data points that are not identified as outliers, which are defined as points beyond 1.5 times the interquartile range.

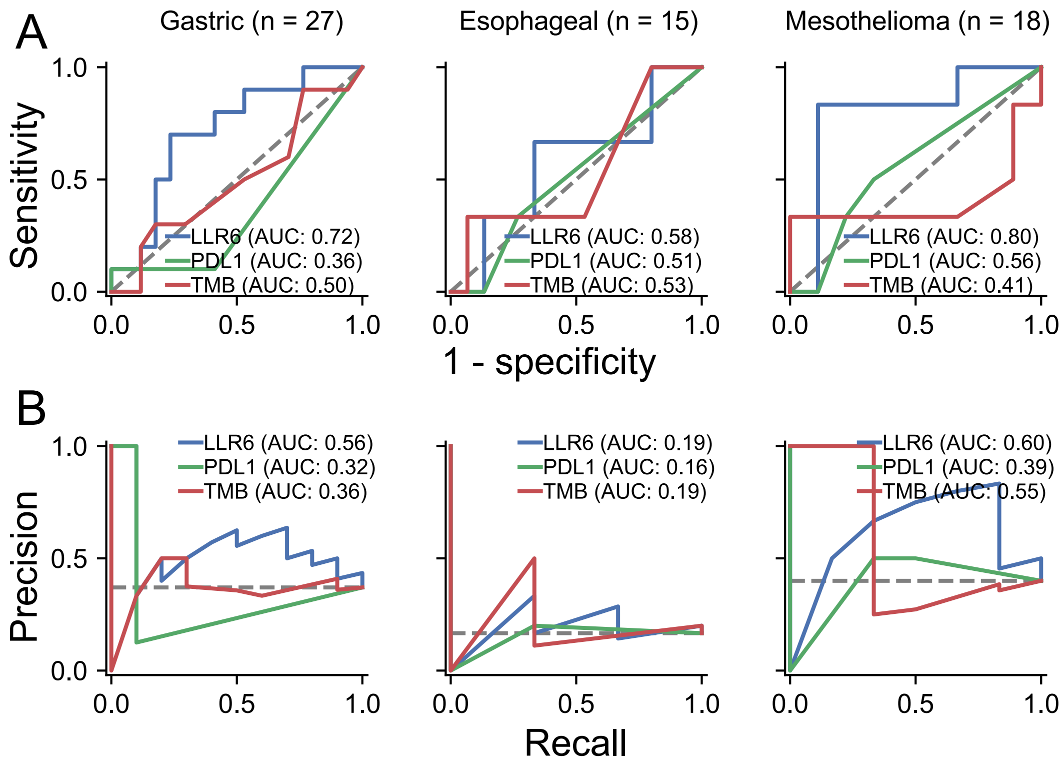


**Supplementary Figure 7. The NSCLC-specific LLR6 model also predicts ICB response in three other cancer types.**

**A.** Receiver operating characteristic curves and corresponding area under the curve (AUC) of the NSCLC-specific LLR6 model, the PD-L1 TPS biomarker, and the TMB biomarker on gastric cancer, esophageal cancer, and mesothelioma.

**B.** Precision recall curves and corresponding area under the curve (AUC) of the NSCLC-specific LLR6 model, the PD-L1 TPS biomarker, and the TMB biomarker on gastric cancer, esophageal cancer, and mesothelioma.
